# Antibodies with Engineered Fc Domains Having Absolute Binding Selectivity to either FcγRIIa or FcγRI Delineate the Respective Effector Phenotypes by Human Monocytes and Macrophages

**DOI:** 10.64898/2026.08.31.748147

**Authors:** Katia George, Sanghwan Ko, Seema Irani, George Delidakis, Chang-Han Lee, Prashant Menon, Jin Eyun Kim, Bart M. Herpers, Evelyn Rocha, Gabriel Soto, Akhil Majmudar, Navin Varadarajan, Y. Jessie Zhang, George Georgiou

**Author notes:** Corresponding Authors: Y. Jessie Zhang and George Georgiou. These authors contributed equally: Katia George and Sanghwan Ko.

## Abstract

IgG1 immune complexes bind to all the Fcγ receptors (FcγR) expressed on myeloid cells, making it challenging to determine the precise role of each FcγR on Fc effector phenotypes. Here we report the engineering of Fc2_KG_, an aglycosylated human IgG1 Fc domain that binds with near physiological affinity to FcγRIIa/b with no detectable binding to any other FcγRs. Crystallographic analysis elucidated the structural basis of how mutations in the Fc domain compensate for the absence of the N297 glycan and enable selective binding. Using single-cell phagocytosis assays, we show that particles opsonized with Fc-engineered antibodies formatted with Fc2_KG_ or with an Fc domain that binds only FcγRI (Fc5), are ingested by THP-1 cells with near identical kinetics. We find that with CD16^+^ primary human monocytes, selective FcγRII engagement is a major contributor to the ADCP mediated by wild-type IgG1. Additionally, we showed that Fc2_KG_ formatted antibodies induce high levels of GM-CSF. With M1-like monocyte-derived human macrophages, trastuzumab formatted with wild-type IgG1 Fc, with Fc2_KG_ or Fc5 were all equally proficient in the trogocytotic killing of opsonized SK-BR-3 HER2^+^ cells but only FcγRI engagement led to secretion of proinflammatory cytokines. Collectively, our results highlight how precisely tuned, Fc-engineered antibodies can be deployed to answer long-standing questions regarding the precise effector functions mediated by human FcγRs, information which is key for the optimization of therapeutic antibodies.

## Introduction

The three protein members of the FcγRII receptor family, FcγRIIa (CD32a), FcγRIIb (CD32b) and FcγRIIc (CD32c) are encoded respectively by FCGR2A, FCGR2B, and FCGR2C (*1*). In primates, FcγRIIa is the most widespread and abundant of all FcγRs and is expressed on Langerhans cells, platelets and on all myeloid cells (*2*). FcγRIIa is an activating immunoreceptor protein, signaling through its own intracellular immunoreceptor tyrosine-based activation motif (ITAM), unlike the other activating Fcγ receptors that rely on the recruitment of the signaling domain FcRγ/FcεRIγ. FcγRIIb is the sole inhibitory receptor and signals through its intracellular immunoreceptor tyrosine-based inhibitory motif (ITIM). It has two isoforms: FcγRIIb1 expressed predominantly on B cells and dendritic cells as well as to a minor degree on monocytes and macrophages (*3, 4*), and FcγRIIb2 which is rapidly internalized upon immune complex binding and is found on myeloid cells, as well as on liver sinusoidal endothelial cells (LSECs), follicular dendritic cells, airway smooth muscle and at low levels on activated T cells (*1, 5*). Finally, FcγRIIc is expressed on NK and B cells only in a subset of the human population (7-15%)(*6, 7*).

The structure and signaling apparatus of FcγRIIa confer this receptor with several unique properties. it is the only FcγR that interacts with all four IgG subclasses(*8, 9*) and the only activating Fc receptor for which antibody binding affinity is largely unaffected by the glycan composition of the receptor or the antibody (*10*). In the human population, there are numerous FcγRIIa SNPs that affect antibody binding affinity (*11, 12*). The most common polymorphism occurs at position 131 (His or Arg)(*11*) with the H131 allelic variant resulting in a slightly higher affinity for IgG1 which has been reported to be of clinical significance in cancer and infectious diseases (*13, 14*).

The engagement of FcγRIIa by IgG opsonized cells or particles can initiate a multiplicity of antibody-mediated effector functions: (1) Phagocytosis-dependent ingestion of bacteria or smaller mammalian cells (*15, 16*). (2) Trogocytosis (by monocytes, macrophages and neutrophils) i.e. the process of uptake of small portions of target cell membrane and associated intracellular material that can in turn result in killing via trogoptosis(*17*). (3) Internalization of immune complexes (*18*). (4) Degranulation of neutrophils which can result in cell killing via ADCC(*19*) and separately, of platelets which can trigger thrombotic events(*20*). (5) Cytokine release (*21, 22*). Finally, (6) there is evidence from murine models that FcγRIIa is critical for the generation of a lasting anti-tumoral memory response following administration of a cytotoxic antibody, a phenomenon termed the “vaccinal effect” (*23*).

Similar to FcγRIIa, the high affinity receptor FcγRI (CD64) is expressed on all myeloid cells (albeit at low levels on resting circulating neutrophils) and it is known to trigger ADCP, trogocytosis/trogoptosis(*24*), degranulation(*25*) and cytokine release (*26*). In blood, cell surface FcγRI is saturated by circulating IgG, leading to its rapid internalization of the receptor followed by recycling to the cell surface independent of ITAM signaling (*27*). However, multivalent immune complexes can displace bound, monovalent IgG from FcγRI leading to immune activation (*28*). FcγRI expression is high in certain tissue resident macrophages, notably at anatomical locations where free IgG concentration is low such as the cervix and alveoli, and is greatly increased on monocyte-derived macrophages under inflammatory conditions (*29–31*).Furthermore, there are reports that under some conditions, engagement of FcγRI may have a significant role on ADCP by tissue resident macrophages even when the expression of the receptor is low (*29*).

A detailed understanding of the relative contribution of the different activating Fc receptors to the IgG effector phenotypes conferred by myeloid cells is important for Fc optimization of therapeutic antibodies to enhance clinical efficacy while minimizing excessive cytokine release. However, clarifying the phenotypes conferred by the individual FcγRs has been challenging for several reasons. As was mentioned above, all myeloid cells express multiple FcγRs at varying levels depending on inflammatory context and anatomical location. The binding of immune complexes by myeloid cells results in the ligation of IgG to all the FcγRs and therefore the ensuing effector phenotypes reflect the integrated response arising from the activation of multiple, distinct albeit overlapping signaling events. There is no facile way to precisely disentangle the contribution of each FcγR to an observed effector phenotype. In principle, this could be accomplished by mechanistic studies whereby all but one Fc receptor on innate cells has been genetically silenced, however this is technically challenging for primary human cells. As an alternative, receptor-blocking F(ab’)₂ fragments or intact anti-FcγR antibodies have been used extensively to isolate the effects of a single Fc receptor while inhibiting others (*11*). Each approach, however, has important limitations. Blocking F(ab’)₂ fragments induce receptor crosslinking, which can perturb membrane protein organization within lipid rafts and thereby affect the localization and signaling of other Fc receptors (*32*). Intact anti-FcγR antibodies present a distinct problem, particularly for FcγRI: rather than occluding the ligand-binding site through their Fab, they act largely through their Fc regionin some cases, actively triggering receptor signaling rather than inhibiting it (*33*). An alternative strategy is to use antibodies having Fc domains engineered to have higher selectivity and affinity towards a certain receptor over others. Beginning with the seminal study by Richards et al.(*34*) numerous combinations of amino acid substitutions and/or glycoengineering have been employed to impart differentially increased FcγRIIa affinity on human IgG1 (*21, 34–43*). However, all the Fc modifications that confer increased affinity and/or selectivity to FcγRIIa reported so far suffer from two limitations: First, engineered Fcs with increased FcγRIIa and/or FcγRIIb affinity also retain varying levels of binding to the other Fc receptors. This is problematic because with high avidity immune complexes even very low affinity binding can result in robust signaling (*37, 43*). Second, nearly all modifications that enhance FcγRIIa/b binding are built on glycosylated IgG1 Fc and the Fc glycans have the potential to engage the two lectin Type II Fc receptors which can affect the function of the Type I or “classical” FcγRs in certain settings (*44*). Two engineered aglycosylated Fc domains that bind to FcγRIIa with affinity even higher than wild-type glycosylated IgG1 have been reported, however both of these variants also bind monovalently to FcγRI with appreciable affinity and, to varying degrees, to FcγRIIIa under high-avidity conditions (*37, 38*).

In this work, we engineered an aglycosylated IgG1 Fc domain (Fc2_KG_) that has absolute selectivity for the FcγRIIa/b receptors. Fc2_KG_ binds to FcγRIIa and FcγRIIb, with K_D_ values nearly identical to that of wild-type glycosylated IgG1, and has no detectable binding to FcγRI, FcγRIIIa, or Type II Fc receptors (since it lacks the N297 glycan) even under very high avidity conditions. The crystal structure of the engineered Fc domain in complex with the clinically relevant FcγRIIa H131 and R131 allotypes helped explain how mutations in the Fc can compensate for the absence of the N297 glycan to confer binding and additionally to impair recognition by FcγRI and FcγRIIIa. Through the use of antibodies formatted with Fc2_KG,_ and separately with an aglycosylated engineered Fc domain that has absolute binding selectivity for FcγRI (Fc5)(*45*), we quantitatively dissected the roles of FcγRI and FcγRIIa on the kinetics and magnitude of phagocytosis by human peripheral monocytes, on trogocytosis by M1-like M(LPS+IFNγ) monocyte-derived macrophages, and on cytokine release by both cell types. We find that the selective engagement of either FcγRI or FcγRIIa alone results in proficient ADCP but very distinct cytokine release profiles depending on cell type.

## Results

### Engineering of FcγRII-selective Fc variants

An IgG1 Fc variant that binds to the FcγRII receptors (FcγRIIa and FcγRIIb) with physiological binding affinity and very highly attenuated binding to FcγRI and FcγRIIIa was first engineered via library screening in *E. coli* using the <u>A</u>nchored <u>P</u>eriplasmic <u>ex</u>pression (APex) display technology (**Fig. 1A**)(*35, 46–48*). In earlier studies, Sazinsky et al.(*37*) had reported that two amino acid substitutions, S298G/T299A, confer binding of aglycosylated IgG1 to FcγRIIa and subsequently Jung et al.(*38*) combined these mutations with amino acid substitutions near the CH2-CH3 interface, to produce Fc1004, which has >100-fold higher affinity towards FcγRIIa_R131_ than wild-type glycosylated IgG1. Both the S298G/T299A and the Fc1004 Fc variants retained significant binding towards the high affinity receptor FcγRI, with K_D_ values in the tens of nM, as well as weak binding to FcγRIIIa. Here we used a combination of site saturation mutagenesis of residues distal to the binding interface coupled with random mutagenesis using Fc1004 as the template. FACS screening for clones displaying antibodies that bind to biotinylated, dimerized FcγRIIa_R131_-GST followed by binding analysis of selected high fluorescence clones on *E. coli* led to the identification of several protein variants which were expressed recombinantly and characterized *in vitro* with respect to FcγR binding properties, eventually leading to the identification of Fc2. Fc2 has very near physiological binding affinity (K_D_) to both the FcγRIIa_H131_ and FcγRIIa_R131_ allotypes as well to FcγRIIb but minimal (i.e. high micromolar, K_D_>10 µM) binding to FcγRI (**Fig. 1B**). Further, Fc2 has very extremely low binding to FcγRIIIa_V158_ (K_D_ = 17 ± 2.35 µM) and no detectable binding by BLI to FcγRIIIa_F158_ or to FcγRIIIb_NA2_ (**Fig. S1A and S1B**). This Fc variant contains six amino acid substitutions: K246Q, T260A, S298G, T299A, A378N and N390D (**Table S1**), of which S298G, T299A and N390D were derived from the Fc1004 template, A378N was introduced by saturation mutagenesis and the remaining two from combinatorial library screening as described above.

**Figure 1.**
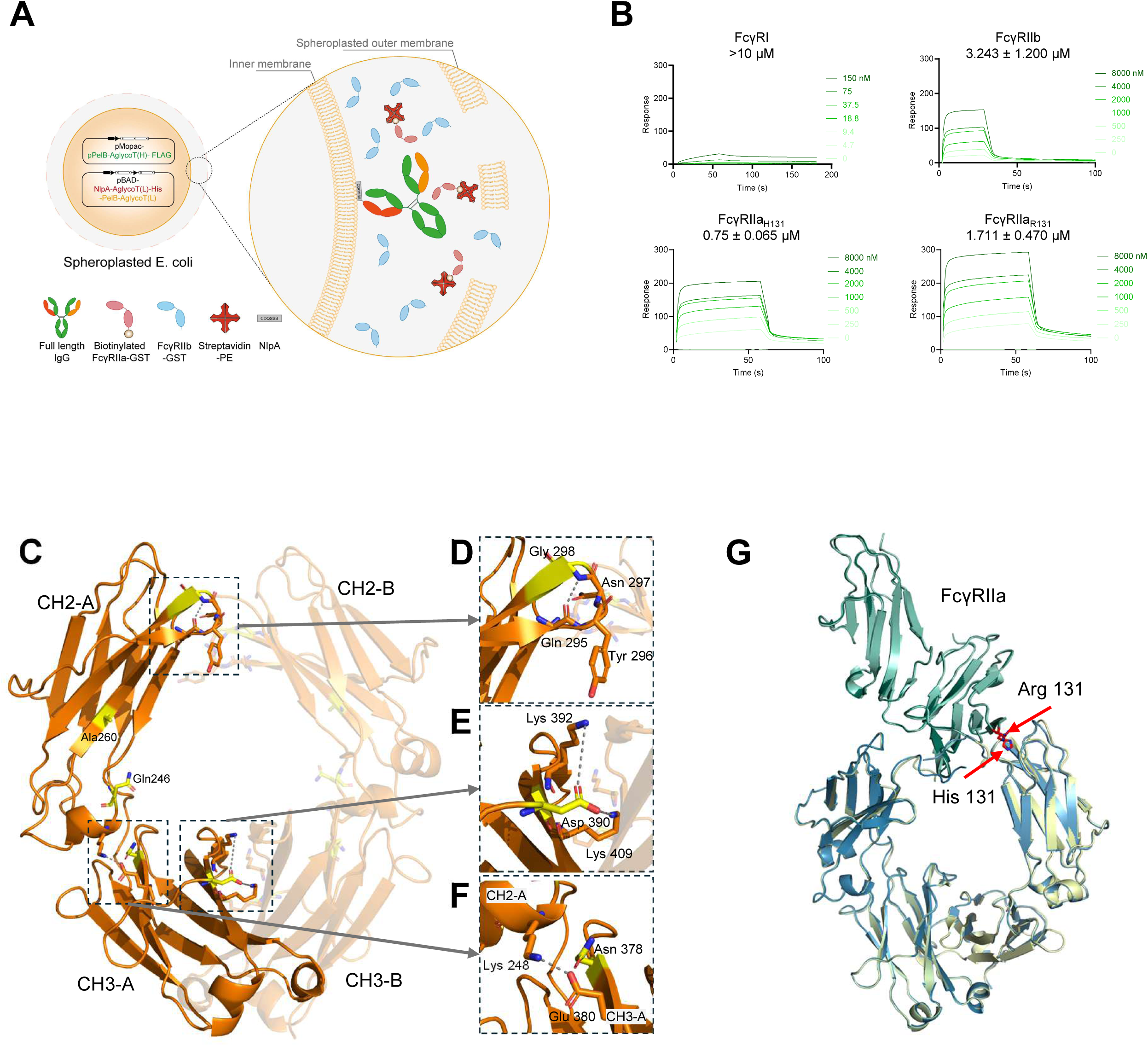
Engineering, FcγR-binding, and structural characterization of Fc2, an engineered IgG1 Fc domain with high selectivity for FcγRII. A) Screening strategy for the isolation of FcγRIIa-selective IgG1 Fc variants by *E. coli* anchored periplasmic display. The VL chain was anchored to the periplasmic side of the inner membrane via an NlpA sequence. Spheroplasts were permeabilized and incubated with unlabeled FcγRIIb-GST and biotinylated FcγRIIa_R131_-GST and streptavidin-PE. B) SPR sensorgrams of Fc2 binding to FcγRI, FcγRIIa_H131_, FcγRIIa_R131,_ or FcγRIIb ectodomain proteins (mean ± SD). C) Crystal structure of apo Fc2 showing the spatial arrangement of the six engineered amino acid substitutions (highlighted in yellow) within the CH2 and CH3 domains of both Fc chains (CH2-A, CH2-B, CH3-A, CH3-B). D) Close-up of the CH2 C’E loop showing Gly298 and Ala299 (S298G/T299A substitutions, yellow). These substitutions promote the formation of a tight β-turn stabilized by a hydrogen bond (dashed line) involving Gly298 and Gln295. E) Close-up of the CH3-A/CH3-B interface showing the N390D substitution which introduces a salt bridge network with Lys392 and Lys409 (dashed lines). F) Close-up of the CH2-A/CH3-A interface. The A378N substitution introduces Asn378, which forms a hydrogen bond with Glu380 (dashed line), which in turn is positioned within salt bridge distance of Lys248, reinforcing CH2–CH3 interdomain coupling. G) Superposition of the Fc2:FcγRIIa_H131_ and Fc2:FcγRIIa_R131_ complex structures. The two receptor allotypes are shown in different shades of green (H131, R131), with the polymorphic residues His131 and Arg131 shown in sticks and indicated with red arrows. The Fc2 chain is shown in yellow and cobalt blue.

### Structure of apo Fc2 and in complex with FcγRIIa_R131_ and FcγRIIa_H131_

The crystal structure of apo Fc2 was determined at 2.6 Å resolution (**Fig. 1C, Table S2**). The overall architecture of apo Fc2 is very similar to earlier published structures of Fc domains bearing a variety of N297 glycans(*49*) (PDB: 3AVE, Cα RMSD= 1.15 Å) or no glycan(*50*) (PDB: 3S7G, Cα RMSD= 1.87 Å) (**Fig. S2A**). The structural analysis of the Fc2 provided a molecular basis for how the mutations in the aglycosylated Fc2 preserved Fc stability in the absence of the glycan, a feature essential for FcγRIIa binding (**Fig. 1C**–**F**). S298G and T299A in the critical CH2 C’E loop are positioned near the Fc–receptor interface, with the Gly and Ala substitutions enabling the formation of a tight β-turn stabilized by a hydrogen bond (Gln295–Gly298) that in turn stabilizes the adjacent structural elements of the receptor binding epitope (**Fig. 1D**). The N390D substitution introduces an acidic residue positioned near Lys392 and Lys409, which may contribute favorable electrostatic interactions that stabilize this region (**Fig. 1E**). In the Fc2 complex, the introduced A378N substitution is positioned near E380. Although the Asn378 and E380 side chains do not form a direct favorable contact, the bulkier Asn378 side chain appears to reposition E380, facilitating the formation of a salt bridge with K248 (2.5 Å) that is absent in wild-type Fc which in turn may be reinforcing CH2–CH3 coupling (**Fig. 1F**). Residue 378 lies within the 370–379 “allosteric band” that spans the CH2–CH3 joint into CH3 and contributes to CH3 stabilization and the CH2–CH3 “ball-and-socket” interface that transmits conformational changes toward the CH2 FcγR-binding surface (*51*). In the native glycosylated IgG1 Fc domain, residues within the 240–260 region contribute to glycan stabilization. Loss of glycosylation in Fc2 removes these interactions, potentially rendering polar or charged residues in this region less favorable. Substitution of K246 with Gln and T260 with Ala likely mitigates this penalty by adapting the local environment in the aglycosylated scaffold to reduce unfavorable interactions which may further assist in the stabilization of the CH2.

We obtained co-crystal structures of Fc2 in complex with FcγRIIa_R131_ and FcγRIIa_H131_ allotypes at 2.7 Å and 3.1 Å, respectively (**Fig. 1G, Table S2**). The two complexes are highly similar (Cα RMSD= 0.4 Å), which is expected given that the two receptor allotypes differ by a single R131H substitution (**Fig. 1G**). The structure of Fc2 in complex with FcγRIIa_R131_ closely resembles that of the wild-type glycosylated IgG1 Fc:FcγRIIa_R131_ complex (PDB: 3RY6, 3.80 Å resolution) with a Cα RMSD of 2.45 Å (**Fig. S2B**). We also confirmed that the aglycosylated Fc2:FcγRIIa_R131_ we report here are highly similar to the higher resolution structures of wild-type glycosylated IgG1 Fc:FcγRIIa_R131_ (PDB: 9MCY, 2.85 Å resolution) reported very recently (Cα RMSD= 1.56 Å) (**Fig. S2C**)(*52*). Receptor binding induces an opening of the CH2-CH2 domain interface — i.e., a separation of the two CH2 domains — compared to the apo Fc2 structure, reflecting the well-established higher conformational flexibility of aglycosylated IgG1 Fc domains (**Fig. S2D and E**)(*50*). Specifically, the distance between the P329 residues of chains A and B increases from 20.1 Å in the apo Fc2 structure to 31.7 Å in the Fc2:FcγRIIa_R131_ complex, i.e. Δ (P329_A_:P329_B_ distance) = 11.6 Å (**Fig. 2D**). The distance between chains A and B in wild-type glycosylated IgG1 Fc:FcγRIIa_R131_ (PDB: 9MCY) is 31.1 Å which is essentially identical to that in the Fc2:FcγRIIa_R131_ complex. We note that this receptor-induced separation of the CH2 domains is much smaller for wild-type glycosylated IgG1 Fc (Δ = 6 Å; compare PDB: 3AVE and 9MCY) than for aglycosylated Fc2 (Δ = 11.6 Å, noted above) (**Fig. S2E**). This convergence of the bound-state geometry suggests that Fc2 preserves the architecture required for productive FcγRIIa engagement in the absence of Fc glycosylation. In the complex, Fc2 adopts the canonical asymmetric binding mode of IgG1 Fc interacting with FcγRs, with chain A engaging the hinge region between domains I and II of FcγRIIa whereas chain B contacts the lateral surface of domain II (*9, 52*). To examine the mode of recognition by the aglycosylated variant, we compared the receptor-contacting loops of Fc2:FcγRIIa_R131_ with those of the recently reported wild-type Fc:FcγRIIa_R131_ complex (9MCY). The lower hinge, BC loop, and FG loop were identical in sequence between Fc2 and wild-type Fc, differing only at S298G in the DE loop (**Fig. 2A**), and were structurally highly similar (**Fig. 2B**). One difference was observed at chain B, where the Fc2 lower hinge sits closer to FcγRIIa than in wild-type Fc (**Fig. 2C–E**). This likely reflects the greater conformational flexibility of the aglycosylated Fc, which permits local adjustment of the lower hinge upon receptor engagement. Thus, despite lacking the N297 glycan, Fc2 engages FcγRIIa through the same structural determinants previously described for glycosylated IgG1 Fc, rather than an alternative or compensatory interface. Collectively, these mutations recapitulate the stabilizing role of the N297 glycan, restoring the architecture required for FcγRIIa engagement.

**Figure 2.**
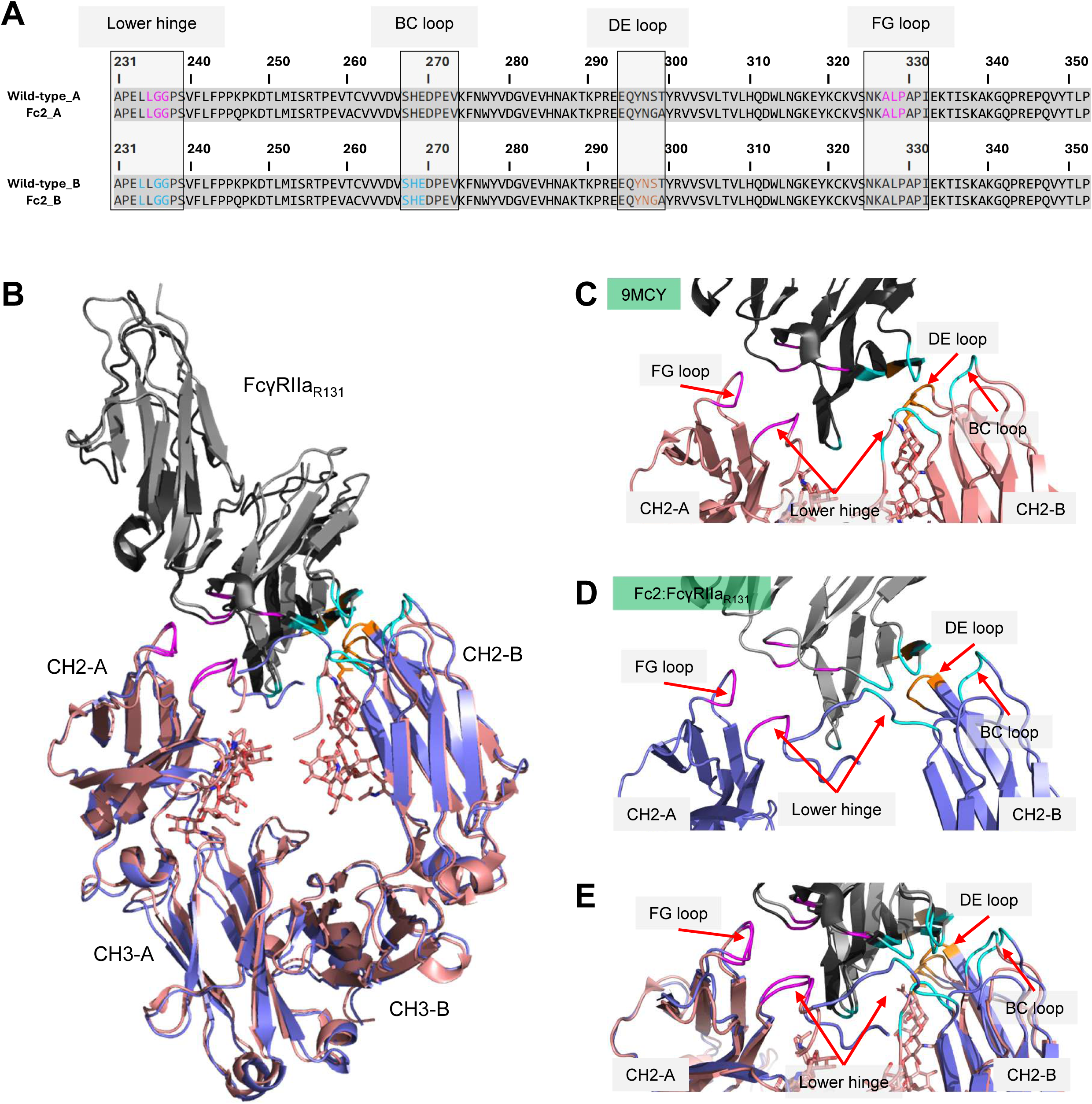
Structural basis for FcγRIIa engagement by the aglycosylated Fc2. **A)** Sequence alignment of wild-type IgG1 Fc and Fc2 (residues 231–350, EU numbering) for chains A and B. The lower hinge, BC loop, DE loop, and FG loop are boxed. **B)** Superposition of the Fc2:FcγRIIa_R131_ complex with the wild-type glycosylated Fc:FcγRIIa_R131_ complex (9MCY). For 9MCY, FcγRIIa is shown in black and the Fc in salmon; for the Fc2 complex, FcγRIIa is shown in gray and the Fc in blue. The two Fc chains are labeled CH2-A/CH3-A and CH2-B/CH3-B. N-glycans are shown as sticks. **C–E)** Close-up views of the Fc–FcγRIIa contact interface. Red arrows indicate the four receptor-contacting loops of the Fc (lower hinge, BC loop, DE loop, and FG loop). Corresponding contact regions on Fc and FcγRIIa are shown in matching colors. Panels show **C)** the wild-type Fc:FcγRIIaR131 complex (9MCY), **D)** the Fc2:FcγRIIaR131 complex, and **E)** the two complexes superimposed.

Separately, the highly attenuated binding of Fc2 to FcγRIIIa (detectable binding by BLI only for FcγRIIIa_V158_ with a K_D_ of 17 ± 2.35 μM compared to 0.113 ± 0.01 μM for wild-type IgG1; **Figs. S1A and S1B**) is likely due to the absence of the N297 glycan, which mediates a direct carbohydrate–carbohydrate contact with the Asn162-linked glycan of FcγRIIIa that is critical for high-affinity binding (**Fig. S3**)(*53*). Unlike the FcγRIIa interface, which Fc2 recovers through protein-intrinsic substitutions, this glycan-dependent contact cannot be compensated by such mutations.

### A G236A substitution in Fc2 confers absolute selectivity towards the FcγRIIa & FcγRIIb

Even though the binding of Fc2-formatted antibodies to FcγRI is highly attenuated with a K_D_ in the 10s of µM, it is nonetheless still measurable by ELISA (**Fig. S4 and Fig. 3B**). Earlier studies had shown that in glycosylated IgG1, a G236A amino acid substitution reduces the K_D_ of glycosylated IgG to FcγRI by 2 to 7-fold (as well as to the two FcγRIIIa allotypes (V158 and F158)) while increasing affinity to FcγRIIa_R131_ and FcγRIIa_H131_ (*34, 43*). Introduction of the G236A substitution into Fc2 completely abolished any residual binding to FcγRI (Fc2 G236A hitherto named Fc2_KG,_ **Table S1, Fig. S4A**). Fc2_KG_ bound to FcγRIIa_R131_ and FcγRIIa_H131_ with an affinity experimentally indistinguishable from that of wild-type IgG (**Fig. 3B**). Additionally, the G236A substitution resulted in a slight (2-fold) reduction in affinity for FcγRIIb for Fc2_KG_ relative to its parental Fc2.

**Figure 3.**
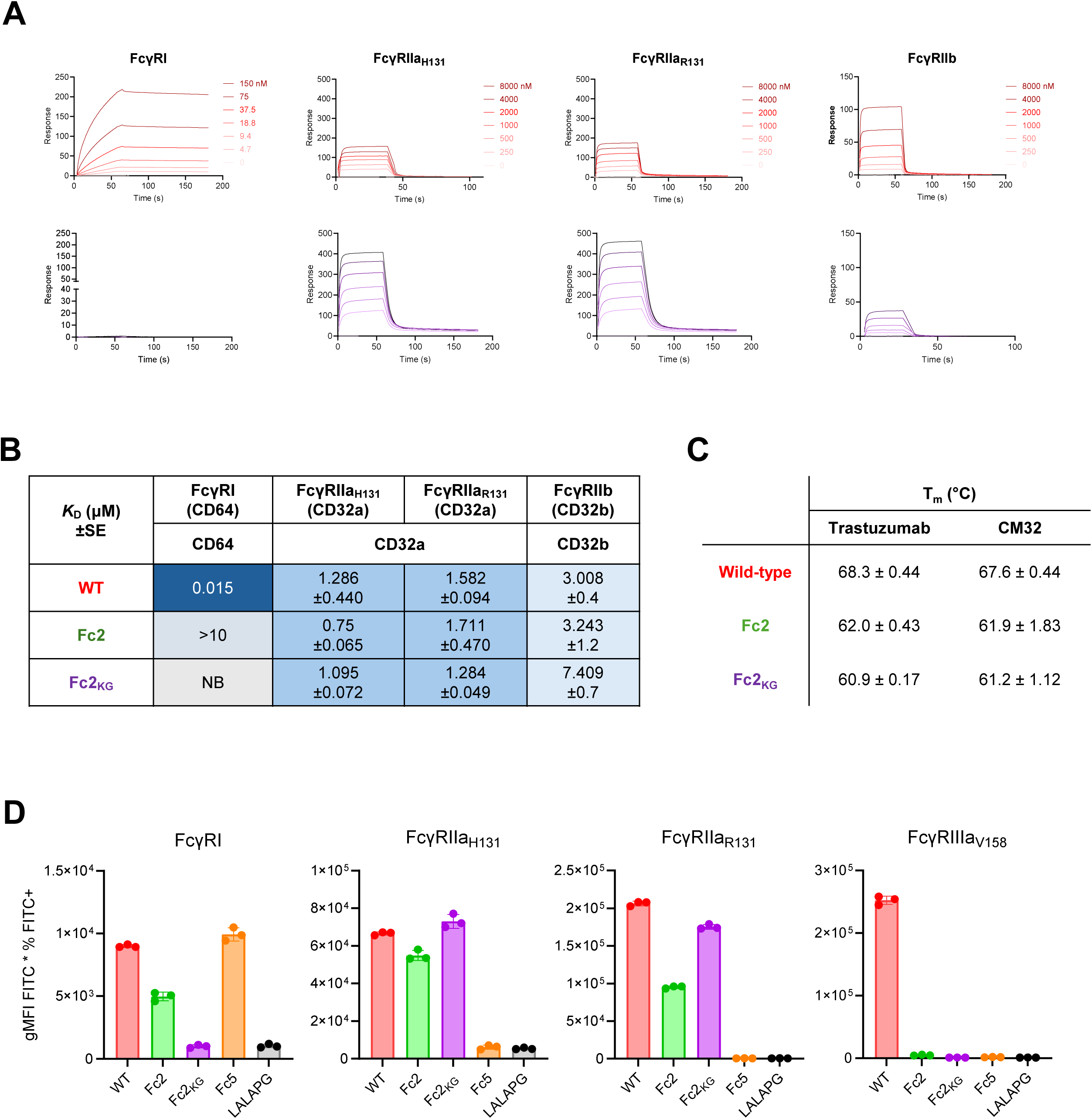
Fc2_KG_ binds exclusively to CD32 receptors with near-physiological affinity. **A**) SPR sensorgrams of CM32-wt IgG1 (above) and CM32-Fc2_KG_ (below), binding to streptavidin-immobilized, biotinylated FcγRs. **B)** Summary of K_D_ values for CM32-wt IgG1, CM32-Fc2, and CM32-Fc2_KG_. The sensorgrams for Fc2 are shown in Figure 1B. All values were calculated using steady-state analysis, except for FcγRI, which was determined using a 1:1 kinetic fit. **C)** Melting temperatures (Tm) of trastuzumab-wt and CM32-wt IgG1, and their Fc2 and Fc2_KG_ variants as determined by differential scanning fluorimetry (DSF). Values represent mean ± SD. **D)** Binding selectivity of immune complexes formed by various trastuzumab variants to CHO cells overexpressing human Fcγ receptors. Immune complexes (250 nM final concentration) were formed by mixing antibody with goat F(ab’)₂ anti-human kappa light chain and incubated with CHO cells stably expressing human FcγRs. Binding was evaluated by flow cytometry using FITC-conjugated AffiniPure rabbit anti-goat IgG F(ab’)₂. Data shown are from a representative experiment with samples run in triplicate.

CM32(*54*) a neutralizing antibody that targets the SARS-CoV-2 Spike protein N-terminal domain and separately, the anti-HER2 antibody trastuzumab(*55*) were formatted with the Fc2 and Fc2_KG_ Fc domains for biophysical and functional studies. For both trastuzumab and CM32, switching the wild-type IgG Fc domain with either the aglycosylated Fc2 or Fc2_KG_ resulted in an approximately 6-7 °C reduction in the Tm as measured by differential scanning fluorimetry (DSF), a reduction in thermal stability which is typical for antibodies lacking the N297 glycan (**Fig. 3C**)(*56*). Nonetheless, the aglycosylated variants retained T_m_ values of ∼61–62 °C, consistent with a stably folded Fc under physiological conditions.

We then tested whether high avidity immune complexes of antibodies formatted with the - Fc2_KG_ domain bind with the expected selectivity to CHO cells stably expressing high levels of the various activating FcγR receptors (**Fig. 3D**). Soluble immune complexes were formed by cross-linking various trastuzumab variants with goat anti-kappa light chain F(ab’)_2_ in a 1:1 molar ratio. Trastuzumab with L234A, L235A, P329G (LALAPG)(*57*) Fc-silencing mutations was used as the negative control. Immune complexes with trastuzumab-Fc2_KG_ had basal levels of binding to FcγRI-expressing CHO cells, identical to trastuzumab-LALAPG whereas immune complexes formed by trastuzumab-Fc5 bound in a manner indistinguishable from that of trastuzumab-wt. Even though the K_D_ of Fc2 to FcγRI is >10 µM (**Fig. 3B**), nonetheless under high avidity conditions we detected appreciable binding of trastuzumab-Fc2 immune complexes to FcγRI-expressing CHO cells. As expected, immune complexes formed with trastuzumab-wt, trastuzumab-Fc2, or trastuzumab-Fc2_KG_ bound CHO cells expressing FcγRIIa_H131_ or FcγRIIa_R131_, whereas trastuzumab-Fc5 and trastuzumab-LALAPG showed basal binding. In contrast, only trastuzumab-wt immune complexes bound FcγRIIIa_V158_-expressing CHO cells. Additionally, rituximab formatted with the Fc2_KG_ or Fc2 Fc domains showed no C1q binding and background levels of complement dependent cytotoxicity with CD20^+^ Ramos cells comparable or lower than those obtained with rituximab-LALAPG (which does not activate complement (**Fig. S4B and S4C**)(*58*).

### Kinetics of FcγRIIa and FcγRI-mediated ADCP by THP-1 monocytic cells

THP-1 is a human monocytic leukemia cell line that is widely used for evaluating ADCP by professional phagocytic myeloid cells (*59*). Consistent with previous reports, we confirmed that THP-1 cells express only FcγRI and FcγRIIa and not FcγRIIb or FcγRIIIa (**Fig. 4A**)(*60*). To evaluate ADCP, 1 μm FITC^+^ and pHrodo-loaded latex beads were conjugated to stabilized trimeric SARS-CoV-2 S protein (SARS-CoV-2 S-6P)(*61*). The pHrodo dye is non-fluorescent at neutral pH and fluoresces only upon acidification within the phagosomal compartment (detected in the PE channel). Fluorescence levels therefore report on the degree of particle internalization by THP-1 cells. The beads were mixed with varying concentrations of CM32 IgG1 Fc variants, and the mixture was incubated with THP-1 cells. Particle internalization into THP-1 cells after 4 hours of incubation was determined by flow cytometry. Cells with beads associated (bound to their surface and/or internalized) are FITC^+^, whereas cells that have executed phagocytosis are FITC^+^ PE^+^ double-positive. Because THP-1 cells can internalize multiple beads in succession, the gMFI (geometric mean fluorescence intensity) of the PE^+^ population is proportional to the number of beads internalized per cell. With glycosylated CM32-wt IgG1, the phagocytic index as a function of antibody concentration showed the characteristic “hook or prozone effect”(*62*) with maximum ADCP observed with 4.44 nM antibody (**Fig. 4B**). At low concentrations, in the range of 0.5-10 nM, the phagocytic index achieved via engagement of FcγRI using CM32-Fc5 immune complexes was slightly higher than that conferred by selective FcγRIIa engagement by CM32-Fc2_KG_ opsonized immune complexes. Within this concentration range, the phagocytic index for particles opsonized with glycosylated CM32-wt IgG1, was approximately equal to the sum of the phagocytic index attained when phagocytosis occurred via FcγRI alone (using CM32-Fc5) and via FcγRIIa alone (particles opsonized with CM32-Fc2_KG_). This indicates that FcγRI and FcγRIIa contribute independently to phagocytosis, and provides functional confirmation that Fc5 and Fc2KG selectively engage their respective receptors. At saturating concentrations of antibody (66.6 nM) the phagocytic index arising from FcγRI engagement by CM32-Fc5 was essentially identical to that achieved with CM32-wt IgG1 and only slightly higher than that achieved via selective FcγRIIa engagement by CM32-Fc2_KG_. We then evaluated phagocytosis in the presence of excess of human serum as a competitor. In the presence of 10% human serum the high affinity FcγRI receptor is saturated by free IgG(*63, 64*) resulting in rapid endocytosis and recycling but without phosphorylation of its ITAM sequence (*26*). Consequently, the phagocytosis of particles opsonized with CM32-Fc5 was completely suppressed whereas that was not the case for particles opsonized with the FcγRII-specific CM32-Fc2_KG_ antibodies. In fact, in the presence of serum CM32-Fc2_KG_ reproducibly resulted in a higher phagocytic index than CM32-wt IgG1 at all but the highest concentration of antibody used (1 µM). Given that CM32-Fc2_KG_ and CM32-wt IgG1 bind to FcγRIIa with essentially the same affinity, the reasons why the former is more efficient in executing ADCP in the presence of competing serum IgG will need to be determined in future studies.

**Figure 4.**
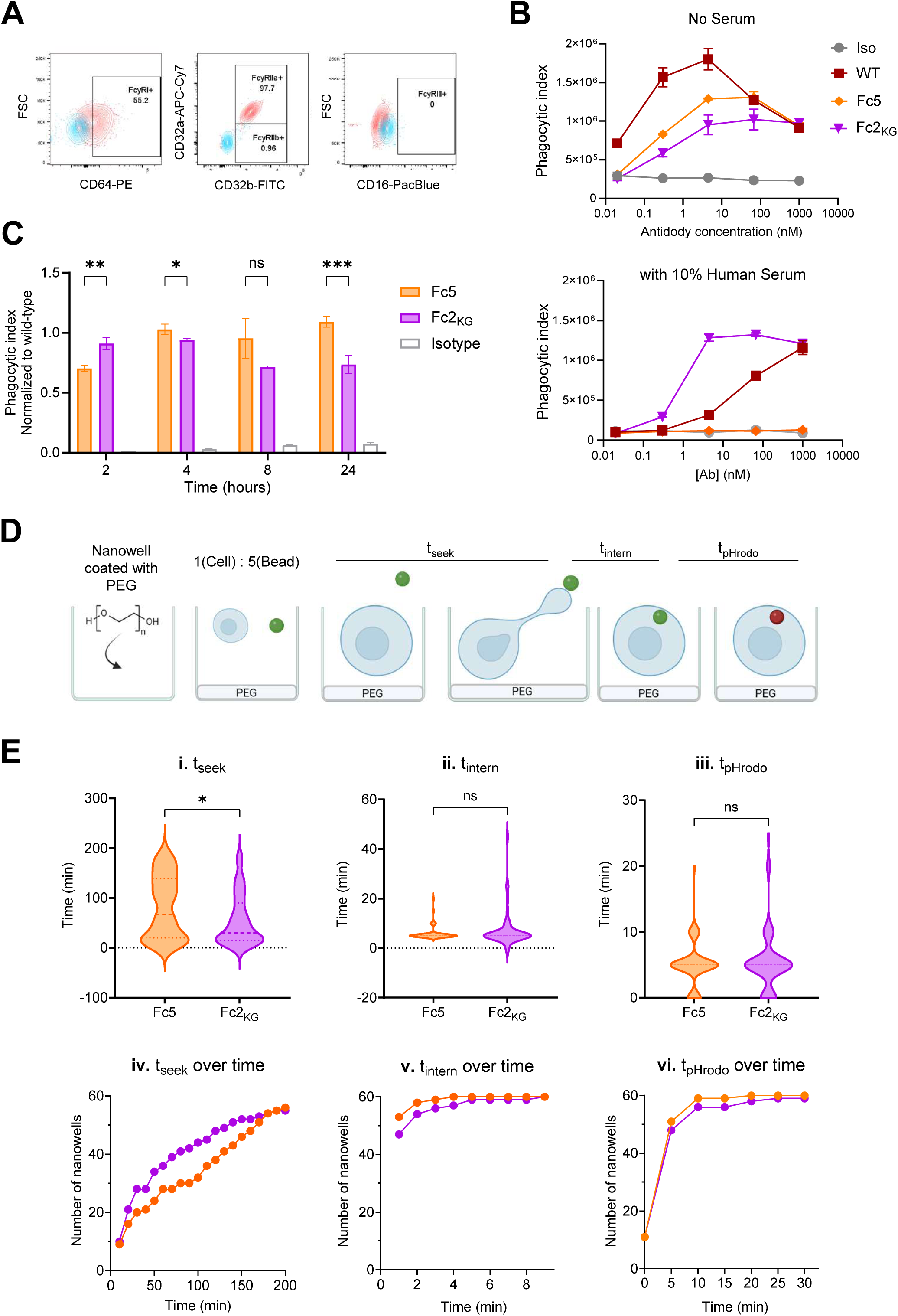
Kinetics of phagocytosis of beads opsonized with CM32-Fc5 or CM32-Fc2_KG_ antibodies with THP-1 cells as effectors. **A)** FcγR surface expression on THP-1 cells determined by flow cytometry using anti-CD64 (FcγRI), anti-CD32a (FcγRIIa), anti-CD32b (FcγRIIb), and anti-CD16 (FcγRIII) antibodies. Percentages denote positive cells within the gate. **B)** THP-1 phagocytic activity as a function of antibody concentration in the absence (upper panel) or presence (lower panel) of 10% human serum. ADCP assays were performed with undifferentiated THP-1 cells and pHrodo-labeled target beads after 4 hours of incubation. Results shown are representative of two independent experiments performed in duplicate. Iso denotes trastuzumab (a non-targeting human IgG1). Error bars indicate ± SD. **C)** Time course of phagocytosis of antibody-opsonized beads by THP-1 cells. THP-1 cells were incubated with HexaPro-coated and pHrodo-labeled DragonGreen beads in the presence of 66.6 nM of the indicated CM32 Fc variants for 2, 4, 8, or 24 hours. Phagocytosis index values were normalized to those obtained with the wild-type IgG1 Fc (CM32-wt IgG1) at each time point. Data shown are the mean of duplicate assays from two independent experiments. Two-way ANOVA: (*) p < 0.05; (**) p < 0.01; (***) p < 0.001; ns, not significant. Error bars indicate ± SD. **D)** Schematic of TIMING assay. Single THP-1 cells within PEG-coated nanowells were incubated with HexaPro-coated and pHrodo-labeled DragonGreen beads coated with 10 μg/mL CM32 antibody variants in a 1:5 cell:bead ratio. **E)** Single-cell time-lapse imaging analysis of ADCP kinetics by the TIMING assay. THP-1 cells were mixed with HexaPro-coated and pHrodo-labeled DragonGreen beads opsonized with Fc5 (orange) or Fc2_KG_ (purple)-formatted CM32 antibody and loaded onto PDMS microwell arrays at a 1:5 cell-to-bead ratio and imaged every 5 minutes for 6 hours. (i–iii) Violin plots showing the distribution of (i) seeking time (t_seek_), the time elapsed from the start of imaging until a cell makes first contact with a bead; (ii) internalization time (t_intern_), the time from first contact until complete bead engulfment; and (iii) pHrodo signal onset time (t_pHrodo_), the time from engulfment until acidification of the phagosome, as indicated by pHrodo fluorescence. Dashed lines indicate median and interquartile range. (iv–vi) Cumulative number of nanowells in which a phagocytic event was completed over time for (iv) bead seeking, (v) internalization, and (vi) pHrodo signal onset. The increasing number of nanowells over time reflects the progressive completion of each phagocytic stage across the imaged cell population. Data represent n = 60 nanowells per condition from a representative experiment. Mann-Whitney test: (*), p<0.05; ns, not significant.

We measured the time course of opsonized particle internalization at saturating conditions (i.e. with 66.6 nM antibody (**Fig. 4C**). Since the purpose of this experiment was to determine the precise contribution of FcγRIIa or FcγRI engagement to ADCP kinetics, the data were normalized relative to the phagocytosis index achieved with glycosylated CM32-wt IgG1. After two hours of incubation, immune complexes formed by CM32-Fc2_KG_ resulted in slightly higher phagocytosis (29.50% ± 10.4%) relative to immune complexes formed by CM32-Fc5. When ADCP was allowed to proceed for a longer time, i.e. 4 hours, CM32-Fc5 conferred a comparable degree of phagocytosis as glycosylated CM32-wt IgG1 and slightly higher than the level achieved with CM32-Fc2_KG_. This was also the case for prolonged phagocytosis experiments i.e. after 24 hr at which point phagocytosis by CM32-Fc5 was 30% higher than CM32-Fc2_KG._ (**Fig. 4C)**.

While the above experiments provide information on the magnitude of phagocytosis for the bulk phagocyte population in the hour time scale, they cannot provide insights on the detailed kinetics of individual particle binding, internalization and endosome transit times, which occur in the minute time scale. To address these questions, we used time-lapse imaging microscopy in nanowell grids (TIMING) (**Fig. 4D and 4E**)(*65*). The time to establish stable conjugation (t_seek_) for ingestion (t_intern_) and for transit to acidified endosomes (t_pHrodo_) were determined for 60 single-cell phagocytosis events per sample. t_seek_ was slightly faster for CM32-Fc2_KG_-opsonized particles relative to particles internalized via FcγRI with the CM32-Fc5 antibody, with the difference reaching statistical significance (**Fig. 4E i and iv**). The faster t_seek_ observed with CM32-Fc2_KG_ relative to CM32-Fc5 may reflect the more rapid association of IgG with FcγRIIa than with FcγRI. However, once a particle became bound to the THP-1 cells, the time constants for ingestion (t_intern_) and for transit to acidified endosomes (t_pHrodo_) were indistinguishable suggesting that the cytoskeletal reorganization events occurring during phagocytosis following engagement of either of the two phagocytic Fc receptors, lead to the transport of opsonized particles to the endosome at the same rate. We note that the faster t_seek_ observed with CM32-Fc2_KG_ immune complexes does not translate to a higher phagocytic index early (**Fig. 4C**) because the phagocytic index reflects not only the kinetics of internalization of individual particles but also the capacity to internalize multiple particles which in turn reflects more complex events related to cytoskeletal rearrangements occurring during repeated cycles of particle internalization.

### Differential roles of FcγRI and FcγRIIa on phagocytosis and cytokine release by peripheral blood monocytes

Human subjects were genotyped for the FcγRIIa_H131_ and FcγRIIa_R131_ allotypes and peripheral blood monocytes were isolated from heterozygous (H/R) donors. The expression of FcγRs on classical (CD14^++^ CD16^-^), intermediate (CD14^++^ CD16^+^) and non-classical monocytes (CD14^low^ CD16^++^) was determined by flow cytometry (**Fig. S5A and S5B)**. Earlier studies have reported that in a fraction of the human population FcγRIIb is expressed above background in both classical and non-classical monocytes (*66*). Hence, to avoid possible confounding effects that might arise from engagement of the FcγRIIb by CM32-Fc2_KG_ antibodies, we selected donors that have background levels of FcγRIIb expression on monocytes. The expression of FcγRI, FcγRIIa and FcγRIIIa in the donors analyzed was consistent with previous reports (**Fig. S5B**)(*66, 67*). ADCP experiments with opsonized beads, after 10 minutes and at later times, ADCP via FcγRIIa (i.e. with CM32-Fc2_KG_ opsonized beads) accounted for progressively a lower fraction of the phagocytic index compared to what was achieved with CM32-wt IgG1. Conversely, the contribution of FcγRI (with CM32-Fc5) increased over time, though it remained below that of FcγRIIa throughout. After 90 minutes, phagocytosis of particles opsonized with CM32-Fc2_KG_ or CM32-Fc5 was still statistically distinguishable but by that time, FcγR-independent particle ingestion of beads opsonized with control antibody (CM32-LALAPG) increased significantly, although remained statistically lower than FcγR-dependent phagocytosis (**Fig. 5A**). Next, we examined the relative contribution of CD16^+^ and CD16^-^ monocytes to phagocytosis after 90 min of co-incubation (**Fig. 5B**). CD16^+^ monocytes are pro-inflammatory, have a more acidified phagolysosomal compartment and are more efficient for ADCP than CD16^-^ monocytes, which however express higher levels of scavenger receptors, and because of their much larger relative numbers in peripheral blood can play a significant role in antibody-mediated phagocytosis in whole blood at least under non-inflammatory conditions (*68–70*). Earlier reports have suggested that FcγRIIIa is critical for tumor cell killing by human monocytes via trogocytosis and ADCC(*71, 72*), but its relative contribution to phagocytosis, compared with FcγRI and FcγRIIa, has been unclear. Although CD16⁺ monocytes (non-classical and intermediate) comprised less than 15% of the total monocyte population (**Fig. S5**), they phagocytosed CM32-wt IgG1-opsonized particles efficiently (**Fig. 5B**), consistent with their established role as potent phagocytic effectors (*66, 67*). With CD16^+^ monocytes, phagocytosis via FcγRI using CM32-Fc5 was very low, though above the isotype control. Given that CM32-Fc2_KG_ accounted for approximately 70% of the wild-type response, the remaining phagocytotic activity of wild-type IgG may reflect a contribution from FcγRIIIa (**Fig. 5B).** Together, these results indicate that phagocytosis by CD16⁺ monocytes is driven predominantly by FcγRIIa, with FcγRI contributing minimally and a possible additional role for FcγRIIIa. Conversely, ADCP by CD16⁻ monocytes was substantially lower than by CD16⁺ monocytes across all antibody formats (**Fig. 5B**).

**Figure 5.**
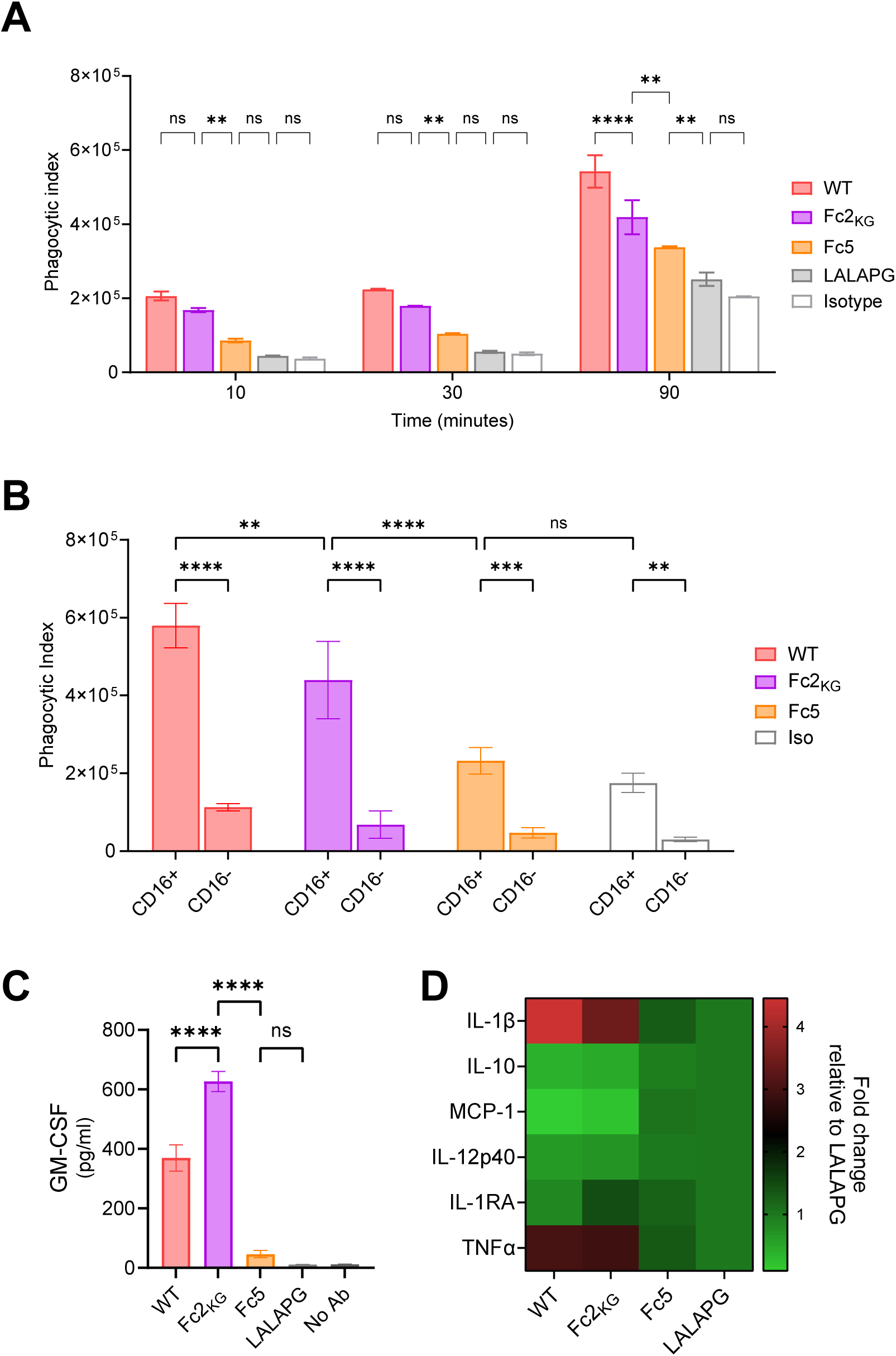
Role of FcγRIIa ligation in ADCP and cytokine release by human peripheral blood monocytes. **A**) Time course of ADCP of opsonized SARS-CoV2 S6P (HexaPro)-coated 1 μm latex beads by classical (CD14⁺CD16⁻) monocytes. Beads were opsonized with 10 nM CM32-wt IgG1, CM32-Fc2_KG_, CM32-Fc5, or CM32-LALAPG and phagocytosis was assessed at 10, 30, and 90 minutes. Data are the mean ± SD of duplicate assays from two FcγRIIa heterozygous (H131/R131) donors. Two-way ANOVA: (*) p < 0.05; (**) p < 0.01; (***) p < 0.001; (****) p < 0.0001; ns, not significant. Error bars show ± SD. **B)** Role of FcγRIIa in ADCP by CD16^+^ and CD16^-^ monocytes. Data shown are the mean of duplicate experiments from each of two FcγRIIa heterozygous H/R donors. Two-way ANOVA: (*) p < 0.05; (**) p < 0.01; (***) p < 0.001; (****) p < 0.0001; ns, not significant. Error bars show ± SD. **C)** GM-CSF levels produced by LPS-activated blood monocytes (n=3 donors) incubated in 96-well plates coated with CM32-wt IgG1, CM32-Fc2_KG_, or CM32-Fc5, CM32-LALAPG, or left uncoated as a negative control. One-way ANOVA: (***) p < 0.001; (****) p < 0.0001; ns, not significant. Error bars show ± SD. **D)** Heatmap summarizing the relative fold-change of cytokine release. Data are reported as fold changes relative to monocytes incubated with the negative control CM32-LALAPG.

Next, we evaluated the role of FcγRIIa and FcγRI engagement in driving cytokine release by LPS-stimulated blood monocytes. Vogelpoel et al.(*73*) reported that LPS-stimulated monocytes secrete cytokines only upon incubation with immobilized IgG or with large immune complexes. Accordingly, we measured cytokine release following incubation of freshly isolated monocytes on IgG-coated plates in the presence of LPS (*73*). For these experiments we used purified antibodies with low endotoxin (<1 EU/mg) and host cell protein levels to preclude confounding effects. Monocytes stimulated with LPS alone secreted modest levels of cytokines while LPS + immobilized wild-type IgG1 resulted in markedly increased secretion (38-fold) of GM-CSF and significant but more modest increases (2 to 3-fold) in the levels of TNF-α and IL-1β(*73*) (**Fig. 5C and 5D**). Unexpectedly, stimulation with antibodies formatted with Fc2_KG_ + LPS resulted in 60% higher GM-CSF secretion relative to wild-type IgG. Stimulation with Fc5 formatted antibodies + LPS led to minimal GM-CSF secretion and no release of other cytokines above the background level detected with LALAPG formatted antibodies.

### The role of FcγRIIa and FcγRI engagement on the phenotypes of M1-like M(LPS+IFNγ) monocyte-derived macrophages

Macrophages kill cancer cells via a combination of phagocytosis and trogocytosis, with trogocytosis possibly being the dominant mode in both 2D and 3D in vitro culture systems especially for high surface density tumor antigens (*74–76*). To investigate the roles of FcγRI and FcγRIIa on cancer cell killing, monocyte-derived M1-like M(LPS+IFNγ) macrophages were differentiated from blood monocytes incubated with M-CSF for seven days followed by activation with LPS and IFNγ (*77, 78*). FcγR expression was evaluated by flow cytometry and was shown to conform to the expected patterns for monocyte-derived M1-like macrophages(*67*) with high levels of expression of FcγRI and FcγRIIa across the population consistent with earlier reports (*26*). 52.3% of the cells expressed FcγRIIb, and a smaller portion (17.8%) expressed FcγRIIIa. M1 polarization was further confirmed by near-uniform surface expression of CD80 (98.3%) (**Fig. S6A and S6B**).

The M(LPS+IFNγ) macrophages were co-incubated with pHrodo-conjugated HER2^+^ SK-BR-3 cells and 5 nM trastuzumab-Fc variants for the degree of target cell material ingestion was quantified by flow cytometry by first gating on CD45 expression, and then for internalization of pHrodo-stained SK-BR-3 cell membrane material in the PE/Cy7 channel. Consistent with earlier findings cited above, live cell confocal imaging revealed that the majority of target cell internalization by macrophages appears to be trogocytotic at least within the time frame of these experiments (**Fig. 6A**). We observed no difference in the time course of target cell material internalization by the effector cells following opsonization with trastuzumab-Fc5, trastuzumab-Fc2_KG_ or glycosylated trastuzumab-wt (**Fig. 6B**). This finding suggests that the selective engagement of either FcγRI or FcγRIIa, or both FcγRs (trastuzumab-wt) leads to the same degree of target cell material internalization.

**Figure 6.**
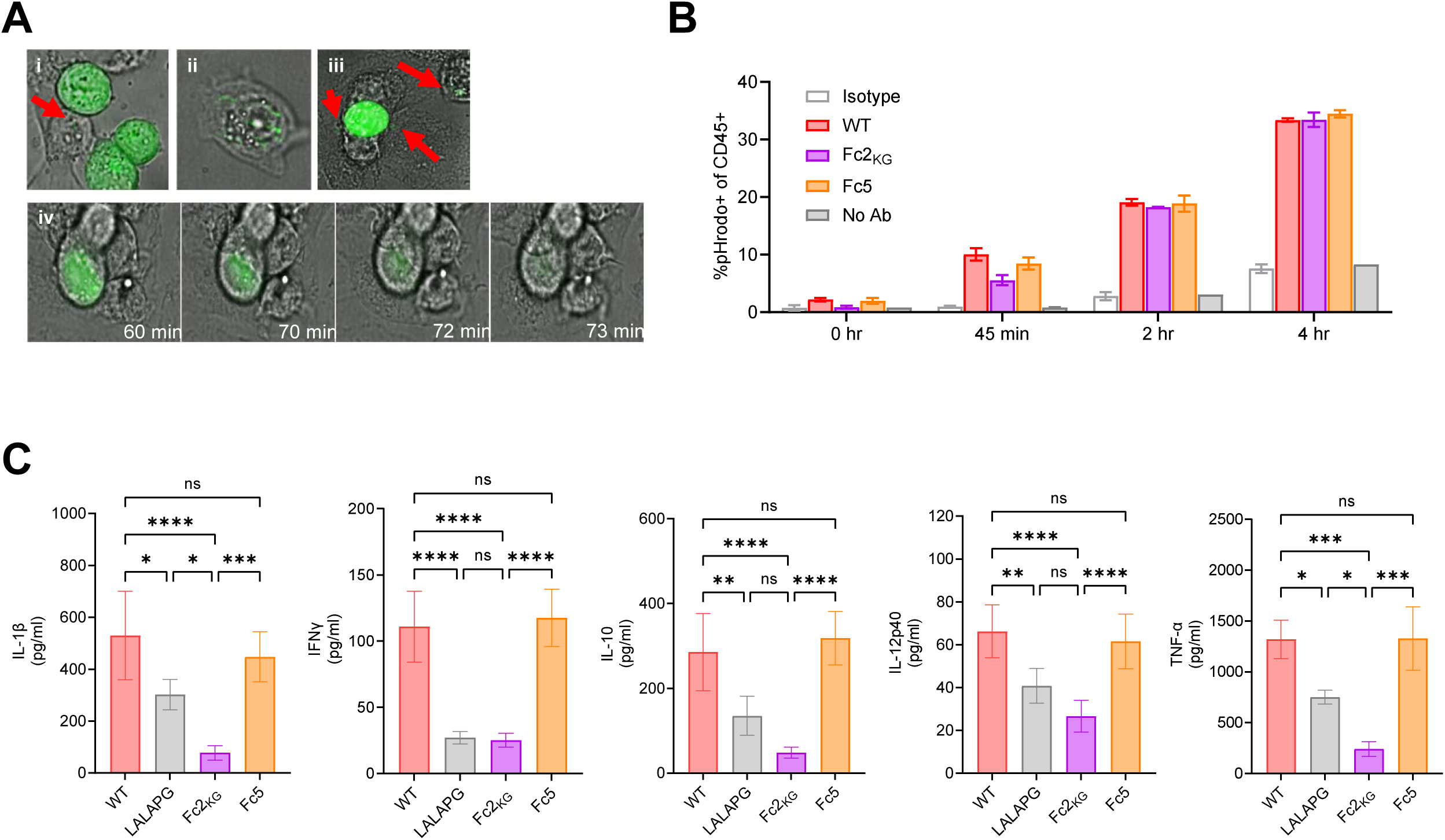
Role of FcγRI and FcγRIIa in the trogocytosis of opsonized SK-BR-3 cancer cells and cytokine release by M1-like M(LPS+IFNγ) macrophages. **A)** Live-cell confocal imaging showing the transfer of calcein-stained SK-BR-3 cell membrane material by M(LPS+IFNγ) macrophages. Macrophages (brightfield) were incubated with calcein-stained SK-BR-3 cells (green fluorescence) and 5 nM soluble trastuzumab-wt. Images were taken every 30 seconds for 2 hours. (i)–(iii), Red arrows indicate macrophages that have internalized calcein-labeled SK-BR-3 membrane. (iv), A representative SK-BR-3 cell positioned above multiple adherent M1 macrophages, showing progressive loss of membrane-associated dye. **B)** Antibody-dependent target cell material internalization by M1 like M(LPS+IFNγ) macrophages. Primary human monocytes were differentiated with M-CSF for 7 days and subsequently polarized with LPS and IFN-γ for 2 days. M1 like M(LPS+IFNγ) macrophages were incubated with pHrodo Deep Red-labeled SK-BR-3 target cells at an E:T ratio of 10:1, in the presence of 5 nM trastuzumab-wt, trastuzumab-LALAPG, trastuzumab-Fc2_KG_, or trastuzumab-Fc5. Macrophages were stained with anti-CD45-PE and the percentage of CD45⁺pHrodo⁺ cells was determined by flow cytometry at the indicated time points. The t=0 hr time point was obtained by immediately detaching macrophages by trypsinization upon mixing. Data shown are from 4 repeat experiments for each of n = 3 donors. Isotype denotes CM32-wt IgG1 (a non-targeting human IgG1), and No Ab denotes the PBS-only (no-antibody) control. Error bars indicate ± SD. **C)** Cytokine release by M1 macrophages upon antibody-dependent stimulation. M(LPS+IFNγ) macrophages were incubated with SK-BR-3 target cells and the indicated trastuzumab Fc variants for 24 hours. Cytokine concentrations in the culture supernatant were determined by multiplex immunoassay. Data shown are from triplicate experiments with cells from one representative donor out of three. One-way ANOVA: (*) p < 0.05; (**) p < 0.01; (***) p < 0.001; (****) p < 0.0001; ns, not significant. Error bars show ± SD.

Finally, we measured the release of cytokines into the supernatant of M(LPS+IFNγ) macrophages that occurred in the course of trogocytosis i.e. as a consequence of co-incubation with SK-BR-3 cells. Trastuzumab-Fc5 antibody induced TNF-α secretion, nearly 2-fold increase over the control (trastuzumab-LALAPG). The level of TNF-α observed with trastuzumab-Fc5 was indistinguishable from that observed with glycosylated trastuzumab-wt. On the other hand, opsonization of SK-BR-3 cells by trastuzumab-Fc2_KG_ failed to induce TNF-α release and in fact it led to an even lower level than SK-BR-3 cells opsonized with trastuzumab-LALAPG (**Fig. 6C**). This may reflect co-engagement of the ITIM-bearing FcγRIIb and/or an inhibitory ITAM (ITAMi) configuration of FcγRIIa, both of which can suppress macrophage pro-inflammatory cytokine secretion (*3, 79*).

Trastuzumab-Fc5 also induced release of IFN-y (4.3-fold over the trastuzumab-LALAPG control), IL-10 (2.4-fold) and a low (approximately 50% increase) change in IL-12p40 and IL-1β. As with TNF-α, the concentration of these cytokines achieved using trastuzumab-Fc5 was indistinguishable from that observed with glycosylated trastuzumab-wt, suggesting that ligation of FcγRI is solely responsible for cytokine release by intact antibodies (**Fig. 6C**). SK-BR-3 cells opsonized with trastuzumab-Fc2_KG_ released lower cytokine levels relative to Fc-attenuated trastuzumab-LALAPG.

## Discussion

While FcγRI, FcγRIIa and FcγRIIIa all signal through ITAM phosphorylation which triggers Syk recruitment and the initiation of the canonical PI3K/PLCγ→NF-κB pathway, differences in affinity for IgG, spatial organization of the respective receptors on the cell membrane and whether signaling depends on the recruitment of FcsRIγ or not are well established to result in distinct signaling biases which in turn lead to overlapping yet distinct phenotypic consequences (*22, 80*). Dissecting the roles of each receptor on myeloid cells which largely express all activating FcγRs, as well as inhibiting receptor FcγRIIb at various levels, has been technically very challenging, yet this information is critical for the optimization of therapeutic antibodies (*15*). We have set out to study this question through the use of engineered aglycosylated IgG1 Fc domains that bind to only to each human FcγR receptor with very high -and in some cases complete selectivity while maintaining physiological affinity and in turn comparing the effector phenotypes with these antibodies relative to the respective wild type, fully glycosylated IgG.

Here we first employed combinatorial library screening strategy involving a combination of random mutagenesis and scanning saturation mutagenesis to engineer Fc2, a human IgG1 Fc variant that binds to FcγRIIa and FcγRIIb with affinities comparable to those of its wild type, glycosylated counterpart. The N297 glycan is critical for the stabilization of the epitope recognized by all Type I FcγRs. While aglycosylated IgG1 Fc domains have been engineered to restore various degrees of FcγR binding (*37, 39, 45, 81*), the structural reasons for how amino acid substitutions compensate for the lack of glycan have not been known. The co-crystal structures of the aglycosylated Fc2 with each of the two major FcγRIIa allotypes (H131 and R131), the first such structures of an engineered aglycosylated Fc bound to an FcγR, reveal that the introduced mutations stabilize the critical C’E loop, which is largely disordered in aglycosylated Fc domains(*50, 82*), and reinforce the CH2–CH3 interface (**Fig. 1C–F**). Together, these observations indicate that the structural role of the N297 glycan can be recapitulated by protein-intrinsic substitutions that rigidify the C’E loop and stabilize the CH2–CH3 interface, providing a structural explanation for how aglycosylated Fc domains recover FcγRIIa and FcγRIIb binding.

These conformational changes likely contribute to the receptor selectivity of Fc2. In particular, the K246Q and T260A substitutions carried by Fc2 may play a role in the pronounced attenuation of FcγRI binding. K246Q removes a positive charge and subtly alters local packing on the CH2 surface, while T260A may tighten local backbone packing and reduce solvent interactions, together potentially affecting shape and charge complementarity at the FcγRI interface(*83*). Interestingly, a thorough recent deep mutational scanning study of glycosylated Fc interactions with FcγR(*51*) revealed that substitutions in each one of residues 246, 260, 378, and 390 were near functionally neutral in terms of FcγRI binding. In contrast, in the context of aglycosylated Fc domains, the substitutions in Fc2 collectively reduce the binding affinity to FcγRI by ∼26-fold (>10 μM for Fc2 vs 382 nM for aglycosylated wild-type IgG1 Fc(*45*)), while the loss of the glycan itself accounts for the larger part of the ∼600-fold reduction relative to glycosylated wild-type IgG1. Structural comparison between Fc domain from the wild-type Fc–FcγRI complex (PDB: 4W4O) and the Fc domain of Fc2 from the Fc2–FcγRIIa_R131_ complex revealed a high degree of structural similarity (Cα RMSD = 1.07 Å), suggesting that the Fc2 substitutions do not significantly perturb the overall Fc fold. Despite the structural resemblance, Fc2 exhibits significantly weaker binding to FcγRI compared to wild-type Fc, implying that local, rather than global, structural differences are responsible. Closer examination revealed a divergence at Fc residue Y296. In the wild-type FcγRI–Fc complex (4W4O), Y296 forms a CH–π interaction with the Cε atom of FcγRI residue K142 (3.4 Å; **Fig. S2F**)(*84*). When the Fc2–FcγRIIaR131 complex is superimposed onto this structure, Y296 adopts an alternative rotamer directed away from K142, beyond the range of a productive contact. This shift likely reflects the altered local environment introduced by the Fc2 substitutions. These observations support the hypothesis that loss of the Y296–K142 CH–π interaction may contribute to the reduced FcγRI affinity of Fc2 (**Fig. S2F**). This structural change, however, did not fully translate into loss of FcγRI binding. Even though the K_D_ value for the binding of Fc2 to FcγRI is reduced by more than 600-fold relative to wild-type IgG1, we found that large immune complexes were still able to bind to CHO cells overexpressing this receptor. In glycosylated Fc domains G236A has been reported to reduce FcγRI binding by ∼7-fold(*34*), and when this mutation was added to the aglycosylated Fc2, it completely abolished binding to this receptor resulting in a selectivity profile that enables binding of immune complexes only to cells expressing FcγRIIa or FcγRIIb (**Fig. 3D**).

Despite extensive efforts, it was not possible to further engineer a variant of Fc2_KG_ capable of binding only to FcγRIIa and not to the highly homologous FcγRIIb (greater than 90% sequence identity, depending on allotype). Thus, Fc2_KG_ enabled us to study pro-inflammatory effector functions mediated by FcγRIIa and not by other activating receptors (in the context of possible inhibitory effects that can result from the co-engagement of FcγRIIb). Using single cell time-lapse imaging analysis, we show that with THP-1 cells as effectors, in the absence of competing IgG, ADCP of opsonized particles via FcγRIIa proceeds with essentially identical kinetics as with FcγRI. These findings suggest that FcγRI may be particularly important for ADCP by monocytic cells in the nasopharyngeal cavity and other tissue locations where extraneous IgG levels are low. However, as had been noted previously(*27, 85*), the high level of circulating IgG in serum results in internalization of FcγRI and blocks ADCP even by high avidity immune complexes.

With primary CD14^+^ monocytes IC internalization via FcγRIIa occurred rapidly and accounted for most of the ADCP in early time points but at later time points phagocytosis of Fc5 opsonized particles via FcγRI engagement increased. By 30 minutes following the onset of the experiment the phagocytic index with Fc5 opsonized particles was only about 25% lower than the value achieved with Fc2_KG_ coated beads. CD16⁺ monocytes are more proficient for ADCP than classical CD16⁻ monocytes. Kang et al.(*71*) demonstrated that CD16 expression contributes to FcγR-mediated phagocytosis in CD16⁺ monocytes, however we show here that engagement of FcγRIIa alone accounts for more than 50% of the phagocytic index attained with fully glycosylated IgG binding to all activating Fc receptors (CD16, CD32 and CD64), indicating that CD32 represents the predominant driver of ADCP even in CD16⁺ monocyte subsets. We further find that FcγRI does not appear to perform ADCP with CD16^+^ monocytes. We note that in studies utilizing FcR blocking antibodies, both FcγRIIIa (CD16) and FcγRI (CD64), but not FcγRIIa (CD32), were shown to mediate antibody-dependent uptake of SARS-CoV-2 into monocytes (*86*). This discrepancy with our findings reported here may be either due to methodological differences, namely the use of blocking antibodies or alternatively, may reflect monocyte activation context-dependent effects. Additionally, we then determined the effect of monocyte stimulation by immobilized Fc2_KG_ or Fc5-formatted antibodies on the release of key cytokines. While earlier studies using blocking antibodies had pointed to FcγRI activation as an important trigger for GM-CSF production by monocytes (*87*), here we found that in fact, only FcγRIIa engaging antibodies were able to induce the release of high levels of GM-CSF as well as other cytokines whereas stimulation by Fc5-formatted antibodies resulted in essentially background levels of cytokine release (**Fig. 5C and 5D**). The differences in effector phenotypes we observe using FcγRI or FcγRIIa selective antibodies relative to earlier studies may reflect the consequences of receptor crosslinking and membrane re-organization arising from the use of blocking antibodies in those reports.

Antibodies targeting highly expressed antigens on cancer cells are known to mediate lethal trogoptosis by myeloid cells, especially with blood-derived macrophages (*75, 88*). M1-like macrophages M(LPS+IFNγ) express high levels of both FcγRI and FcγRIIa. With SK-BR-3 breast cancer cells that express high level of HER2 as targets, we observed that both trastuzumab-Fc5 and trastuzumab-Fc2_KG_ executed trogocytosis with the same kinetics and extent as glycosylated trastuzumab-wt. In other words, engagement of both receptors which occurs by trastuzumab did not lead to enhanced trogocytosis, suggesting that under these conditions the proficiency of M(LPS+IFNγ) to ingest target cell material is not affected by which or how many activating FcγR are involved but rather it is dictated by other cellular processes in endocytosis. We show that under these experimental conditions at least, cytokine release by M1-like macrophages was solely mediated by FcγRI, consistent with earlier findings by Brezski and coworkers (*21*). That latter study also showed that FcγRIIIa can also drive cytokine release by PBMCs.

## Methods

### Human Samples, Cell Lines, and Reagents

SK-BR-3 cell line (ATCC® HTB-30™), Ramos (CRL-1596™), and THP-1 cells (TIB-202^TM^) were obtained from American Type Culture Collection (ATCC). SK-BR-3 cells were cultured in complete Dulbecco’s modified Eagle’s medium (DMEM) with 10% fetal bovine serum (FBS). Ramos and THP-1 cells were cultured in complete RPMI 1640 with 10% FBS. Whole blood was collected from anonymous healthy donors in EDTA-coated vacutainers (Versiti Blood Center). Human monocytes were purified using histopaque density gradient medium (Sigma Aldrich) and the RosetteSep™ Human Monocyte Enrichment Cocktail (STEMCELL), according to manufacturer’s instructions. For the generation of M1 macrophages, monocytes were cultured in RPMI 1640 with 20% FBS and 100 ng/ml M-CSF (Biolegend) at a density of 1 × 10^5^ cells/cm^2^ for 7 days, followed by 2 days of polarization with 100 ng/ml LPS (eBioscience) and 20 ng/ml IFNγ (Biolegend).

### Preparation of recombinant proteins

Recombinant proteins were generated and purified as previously described (*48*). In brief, human Fc receptors (FcγRI, FcγRIIaH/R, FcγRIIb, FcγRIIIaV/F, FcγRIIIbNA1/NA2) and HexaPro were cloned into the mammalian expression vector pcDNA3.4 (Invitrogen) by Gibson assembly. Engineered Fc domains were cloned in-frame into the same vector containing the VH1-CH1 domains of the desired antibody format (i.e., trastuzumab, rituximab, CM32). Proteins were transiently transfected into Expi293F cells (ThermoFisher) using Expifectamine (ThermoFisher). Antibody heavy chain plasmids were transfected with a 3:2 light chain: heavy chain molar ratio. Following 5 days of incubation at 37°C with 8% CO_2_, proteins were purified from the culture supernatant. Antibodies or His-tagged Fc receptors and HexaPro were purified using Protein G Plus affinity chromatography or Ni-NTA affinity chromatography, respectively. Proteins were eluted using 100 mM glycine pH 2.7 or 300 mM imidazole in PBS, followed by a buffer exchange to PBS using Amicon® Ultra Centrifugal Filters, 10 kDa (Millipore). The purity of recombinant proteins was confirmed to be over 95% using SDS-PAGE and any antibody dimers were removed via size exclusion chromatography. When relevant, Fc receptors containing a C-terminal 6 × His-tag and an N-terminal AviTag™ were enzymatically biotinylated using the biotin ligase enzyme, BirA (Sigma Aldrich).

### T_m_ measurement by Differential scanning fluorimetry (DSF)

Purified antibodies at concentrations ranging from 0.25-1.0 mg/ml were mixed with SYPRO Orange Dye (ThermoFisher) according to manufacturer’s instructions. DSF was performed using ViiA 7 Real-Time PCR system (Applied Biosystems) with FAM filters and HID Real-Time PCR Analysis Software v1.3 (ThermoFisher).

### Library construction, display and screening

*E. coli* libraries expressing full antibodies on the inner membrane of E. coli were constructed using a two-plasmid system, as previously detailed (*35, 38, 89*). Briefly, libraries of Fc variants were constructed using appropriate degenerate primers, joined to trastuzumab Fab-CH1 fragment via overlap-extension PCR and the product was ligated into pPelB-AglycoT(H)-FLAG vector (*38*). The DNA was transformed into *E. coli* Jude1 containing the pBAD-AglycoT(L)-His which encodes NlpA-fused trastuzumab light chain for inner membrane anchoring of the IgG1. Transformants were diluted into TB media, cultured for 2 hours at 37°C with shaking and protein expression was then induced by adding 1 mM isopropyl-1-thio-β-D-galactopyranoside (IPTG, Fisher) and 2% (v/v) D-arabinose. After overnight incubation at 25°C the cells were subjected to enzymatic and osmotic treatment to remove the outer membrane, resulting in spheroplasts. Spheroplasts in turn were stained with biotinylated GST-tagged receptors. Binding was detected using streptavidin-PE (Biolegend) and positive events were sorted on a FACS Aria II (BD Biosciences). The Fc genes from sorted spheroplasts were amplified by PCR and re-cloned into the pPelB-AglycoT(H)-FLAG vector for subsequent rounds of sorting and enrichment.

### ELISA analysis

A high-binding 96-well plate (Corning 3361) was coated with 0.5 µg of antibody overnight at 4°C. The plate was blocked with 3% BSA in PBS for 90 minutes at room temperature. Biotinylated and His-tagged FcγRs were tetramerized by adding 100 nM streptavidin to 400 nM receptor. Tetramerized receptors were serially diluted in 3% BSA and incubated on the plate for 1 hour shaking at room temperature. Following a wash with PBST, the plate was incubated with 1:5000 diluted HRP-anti-His Tag secondary antibody (Biolegend) for 30 minutes shaking at room temperature. To measure affinity to C1q, monomeric human C1q protein (Fisher Scientific) was added instead of tetramerized FcγR and binding was detected using a 1:200 diluted HRP-anti-C1q secondary antibody (Fisher Scientific). The plate was then developed with TMB substrate (ThermoFisher) and quenched by 2 M H_2_SO_4_. The absorbance at 450 nm was measured using the Synergy H1 plate reader by BioTek.

### Affinity measurement by surface plasmon resonance (SPR)

SPR was performed on a Biacore X-100 instrument and data analysis was performed using the BIAevaluation software. Biotinylated receptors were immobilized on a CAP sensor chip using the Biotin CAPture kit (Cytiva Biosciences), according to manufacturer instructions. CM32 IgG1-Fc variants were serially diluted in 1X HBS-EP^+^ buffer (0.01 M HEPES, 0.15 M NaCl, 0.005% Tween 20, 3 mM EDTA) at pH 7.4. 8000-250 nM of antibody was used for affinity measurements towards low/medium affinity receptors (FcγRIIa_H131_, FcγRIIa_R131_, and FcγRIIb) and 150-4.7 nM for FcγRI. All antibodies were allowed to associate and dissociate for 60 seconds and 120 seconds, respectively. The chip was regenerated after each binding interaction using the buffer provided by the manufacturer. The K_D_ was calculated using a steady state analysis.

### Affinity measurement with biolayer interferometry (BLI)

The binding affinities of Fc variants towards FcγRIIIa and FcγRIIIb was measured using an Octet RED96 System (ForteBio). Biotinylated receptors and rituximab-formatted antibodies were diluted in Octet Kinetic Buffer (PBS + 0.02% Tween20, 0.1% BSA, and 0.05% sodium azide). Receptors were loaded onto High Precision Streptavidin (SAX) biosensors to a response of 0.5 nm. Serially diluted antibodies were then associated for 2 min and dissociated for 2 min while shaking at 1000 rpm at 25°C. Biosensors were regenerated for 30 seconds in 2 M MgCl_2_. K_D_ values were calculated using a steady state analysis in BIAevaluation software. Sensorgrams were acquired using multi-cycle kinetics, with each concentration measured as an independent cycle and regeneration between injections. Individual cycles were compiled into a single panel for visualization. Baseline correction was applied to each cycle prior to K_D_ determination.

### Crystallization of Fc2 and Fc2 in complex with FcγRIIa

Fc2 crystals were obtained by seeding crystals into drops from the mother liquor containing 15% to 22% PEG 3350 pH 5.0 to 6.5. Crystals were then vitrified in liquid nitrogen after a brief soaking in the mother liquor containing 15% glycerol. Crystals diffracting to a resolution of 2.6 Å were obtained at the APS beamline 23-ID-D. The Fc2:FcγRIIa complexes were generated by incubating excess Fc2 with either FcγRIIa_H131_ or FcγRIIa_R131_ at 5 mg/ml and molar ratio of 1.2 for 1 hour at 4°C. Complexes were purified on a Superdex^®^ 200 Increase 10/300 GL (GE Healthcare) size-exclusion column and subsequently concentrated to 8mg/ml using a 10 kDa Vivaspin^®^ Turbo ultrafiltration concentration (Sartorius AG). Crystallization screening trays were set up using an Art Robbins crystallization robot. Needle-shaped crystal hits were obtained in the screening trays with 0.1 M HEPES pH 7.0, 0.1M KCl, 15% PEG 5000 MME, and were subsequently optimized in 24-well trays using sitting drop vapor diffusion at 4°C. Crystals were briefly soaked in a solution of mother liquor with 15% glycerol before vitrification in liquid nitrogen. FcγRIIa_R131_ complexes that generated crystals with low diffraction were obtained in a 24-well plate with 12% PEG 5,000 MME, 0.1M HEPES pH 7.0 and 0.1 M KCl and were used to seed larger and better diffracting crystals in 12-17% PEG 5,000 MME within pH range 6.75 to 7.25. Complexes were subsequently vitrified in a cryo chamber containing the mother liquor and 15% glycerol to produce diffraction data at 2.7 Å at the Advanced Light Source (ALS) BL 5.0.1. Crystals of FcγRIIa_H131_ complexes diffracting to a resolution of 3.1 Å at Advanced Photon Source (APS) BL 23-ID-D source were formed in mother liquor of 12% PEG 5000MME 0.1M HEPES pH 7.0 and 0.1 M KCl and vitrified in mother liquor containing 15% glycerol. The structures of Fc2 in complex with the FcγRIIa_H131_ and _R131_ alleles were solved by molecular replacement using Phaser on Phenix suite(*90*) with the PDB structure of 3WJJ(*91*) as a search model and that of Fc2 was obtained using 3AVE as a search model. The refinement was conducted using the refinement program in the Phenix suite (*92, 93*). Model building of sugar molecules was done manually on COOT (*94*).

### High avidity binding characterization using CHO-K1 cells

Immune complexes were generated by incubating Herceptin formatted Fc variants in a 1:1 molar ratio with Goat F(ab’)_2_ anti-Human Kappa Light Chain (Bio-Rad) for two hours at 37°C with gentle rotation and then placed on ice. CHO-K1 cells stably transfected with either FcγRIIa_H131_ (ECACC# CHO-K1.Cl-0205), FcγRIIa_R131_ (ECACC# CHO-K1.Cl-0204), FcγRIIIa_V158_, (ECACC# CHO-K1.Cl6), or FcγRI (Acro Biosystems# SCCHO-ATP062L), were thawed, washed, and plated in a 96-well tissue cultured plate at 100,000 cells per well and then mixed with 100 μl of immune complexes at a final concentration of 250 nM and incubated on ice for 30 minutes. The cells were subsequently washed twice in FACS buffer and resuspended in (FITC)-AffiniPure F(ab’)_2_ Fragment Rabbit Anti-Goat IgG, F(ab’)_2_ Fragment Specific (Jackson ImmunoResearch) at 10 μg/ml diluted in FACS buffer and incubated on ice for 30 minutes. Cells were washed twice in FACS buffer and immune complex binding was determined on a BD Fortessa at the Center for Biomedical Research Support Microscopy and Imaging Facility at UT Austin (RRID:SCR_021756). Flow Cytometry data was analyzed using FlowJo™ v10.8.1 (BD^®^).

### FcγR Polymorphism Genotyping

Donor gDNA was isolated using the DNeasy Blood and Tissue Kit (Qiagen) and genotyped using the Applied Biosystems^®^ ViiA^™^7 Real-Time PCR System. The following TaqMan^®^ SNP Genotyping Assays were used to genotype FCGR3A and FCGR2A, respectively: Assay ID C_25815666_10 (SNP ID: rs396991), and C_9077561_20 (SNP ID: rs1801274).

### Flow Cytometry

Cells were prepped for flow cytometry with several washes in FACS buffer (PBS with 2% FBS and 5mM EDTA) followed by 30 minutes staining at 4°C with the following antibodies (Biolegend) at 1:200 dilution, as applicable: anti-CD14-AF488 (63D3, Cat # 367130), anti-CD16-Pacific Blue^™^ (3G8, Cat #980106), anti-CD32a-APC/Fire^™^750 (FUN-2, Cat #303220), anti-CD32ab-FITC (IV.3, STEMCELL Cat #60012Fl.1), anti-CD32b-AF647 (S18005H, Cat #398306), anti-CD64-PE (10.1, Cat #305008), anti-CD45-PE (HI30, Cat #982322). Flow cytometry was performed using the BD Fortessa and data was analyzed using FlowJo^™^ v10.8.1 (BD^®^).

### ADCP Assay with THP-1 cells and Primary Monocytes

Antigen-coated beads were generated by incubating 1 μm DragonGreen beads (5 × 10^8^ bead) (Bangs Laboratories) with 25 μg of the HexaPro variant of the SARS-CoV-2 S protein(*54*) for 1 hour rotating at room temperature. In parallel, ultra-low IgG FBS (Gibco) at 5% in PBS was dyed with pHrodo^™^ iFL Red STP Ester (amine-reactive) (ThermoFisher) for 1 hour at room temperature, according to manufacturer instructions. HexaPro-coated beads were then blocked with pHrodo^™^-labeled FBS for 1 hour rotating at room temperature to generate pHrodo-labeled, HexaPro-coated beads. Following a wash using Costar^®^ Spin-X centrifuge tubes with a 0.22 μm pore filter (Corning), immune complexes were reconstituted in 5% ultra-low IgG FBS in PBS. Immune complexes were then co-incubated with THP-1 cells in a 1:100 cell:bead ratio in a 96-well plate, alongside indicated CM32-Fc variant concentrations in RPMI 1640 for indicated duration at 37 °C – either with or without 10% human AB blood type serum (Sigma Aldrich). Plates were then washed with FACS buffer by centrifuging at 300 × g for 10 minutes, resuspended in FACS buffer and analyzed via flow cytometry.

Primary monocyte ADCP assays were performed by incubating freshly isolated cells with immune complexes at 1:20. Prior to FACS analysis, monocytes were labelled with anti-CD16– Pacific Blue antibody to identify the monocyte population by flow cytometry. All steps, including immune complex generation, were performed in sterile conditions using a biosafety cabinet.

### Trogocytosis/phagocytosis assays with M1-like M(LPS+IFNγ) Macrophages

SK-BR-3 cells were labeled with the pHrodo^™^ Deep Red Mammalian and Bacterial Cell Labeling Kit (ThermoFisher) according to manufacturer’s instructions. For the generation of M1-like polarized macrophages, (M(LPS+IFNγ) macrophages) primary human monocytes were cultured in RPMI 1640 with 20% FBS and 100 ng/ml M-CSF (Biolegend) at a density of 1 × 10^5^ cells/cm^2^ for 7 days, followed by 2 days of polarization with 100 ng/ml LPS (eBioscience) and 20 ng/ml IFNγ (Biolegend). M1-like polarized macrophages in 24-well TC-treated plates (Corning) at a density of 2 × 10^5^ cells/well were centrifuged in their plate at 700 × g for 5 minutes to encourage plate adherence. Supernatant was aspirated and 2 × 10^4^ SK-BR-3 cells were added alongside diluted anti-HER2 (Trastuzumab). After incubation at 37°C for the indicated time, media containing suspended cells was collected and adherent macrophages were detached with 0.25% Trypsin-EDTA (Fisher Scientific) and then recombined with the previously collected media in a non-TC treated 96-well deep-well plate (Corning). When applicable, FcγRIIb on M1 macrophages was blocked by incubating cells with 10 μg/ml of a variant of the anti-FcγRIIb antibody 2B6^57^ containing the LALAPG mutations for 15 minutes, at 37°C prior to the addition of SK-BR-3 cells. Cells were washed with FACS buffer by centrifugation at 600 × g for 10 minutes and then stained with anti-CD45-PE to label M1 macrophages. Trogocytosis was determined based on the percentage of CD45^+^ cells that were also pHrodo^™^ positive. All steps were performed in sterile conditions using a biosafety cabinet.

### TIMING Assay

TIMING experiments were performed as previously described (*65*). Briefly, HexaPro-coated and pHrodo-labeled 1 μm DragonGreen beads were generated as described above and incubated with appropriate antibody at 10μg/ml at room temperature for 1 hour. Unstained THP-1 cells were loaded onto a PDMS microwell array alongside the coated beads at a 1:5 density. The microchip was imaged in three channels: brightfield, FITC/DragonGreen (exposure time: 60 ms), and TexasRED/pHrodo (exposure time: 200 ms). Images were taken every 5 minutes for 6 hours.

### Cytokine Release Assay

96-well flat-bottom plate (Corning) were coated with 100 ng of appropriate antibody variant overnight at 4°C, the plates were washed in PBS and blocked with 2% FBS in PBS for 2 hours at 37°C. Monocytes were seeded in the mAb-coated plate at a density of 50,000 cells per well in RPMI 1640 with 10 ng/ml LPS. 24 hours after incubation at 37°C, the supernatant was collected and cytokine levels were determined as described below. Similarly, the supernatants from 24-hour trogocytosis assays with activated, M1-polarized monocyte-derived macrophages as effectors and SK-BR-3 cells as targets were collected and analyzed. Cytokine levels in culture supernatants were determined using the TNF-α ELISA MAX^™^ Deluxe ELISA Kit (Biolegend) according to manufacturer instructions and the Human Cytokine Proinflammatory Focused 15-Plex Discovery Assay^®^ Array (HDF15) by Eve Technologies (Calgary, AB).

### Live Cell Confocal Imaging

SK-BR-3 cells were stained with calcein AM fluorescent dye (ThermoFisher) according to manufacturer instructions, then co-incubated with 5nM WT Trastuzumab and M1 macrophages cultured on a 6-well plate containing a No. 1.5 uncoated coverslip with a 14mm glass diameter (MatTek Corporation) at a 1:10 ratio. Images were captured on a Nikon CSU-W1 spinning disk confocal on a Nikon Ti-2 inverted microscope equipped with stage top incubation and a Nikon Plan Apo VC 60x/1.2 water immersion objective. Fluorescence was excited with a 488 nm laser and collected with a 525/36 emission filter. Images were captured using an Andor iXon Ultra 888 EMCCD camera controlled with NIS-Elements software. Time-lapse fluorescence and brightfield images were collected every 30 seconds for 1 hour, using the Nikon Perfect Focus System to maintain the focal plane. Microscopy was performed at the Center for Biomedical Research Support Microscopy and Imaging Facility at UT Austin (RRID:SCR_021756).

### Data and materials availability

Atomic coordinates and structure factors for the reported crystal structures have been deposited in the Protein Data Bank under accession codes 36JH (apo Fc2), 36JI (Fc2–FcγRIIa_R131_ complex), and 36JJ (Fc2–FcγRIIa_H131_ complex).

### Data Analysis

Data were analyzed for statistical significance using GraphPad Prism 9 (GraphPad Software).

## Acknowledgments

We thank Jack Borrok for their help with advice on structural analysis.

## Table of Contents for Supplementary Materials

Figures S1 to S6

Table S1 and S2

---

**Figure S1.**
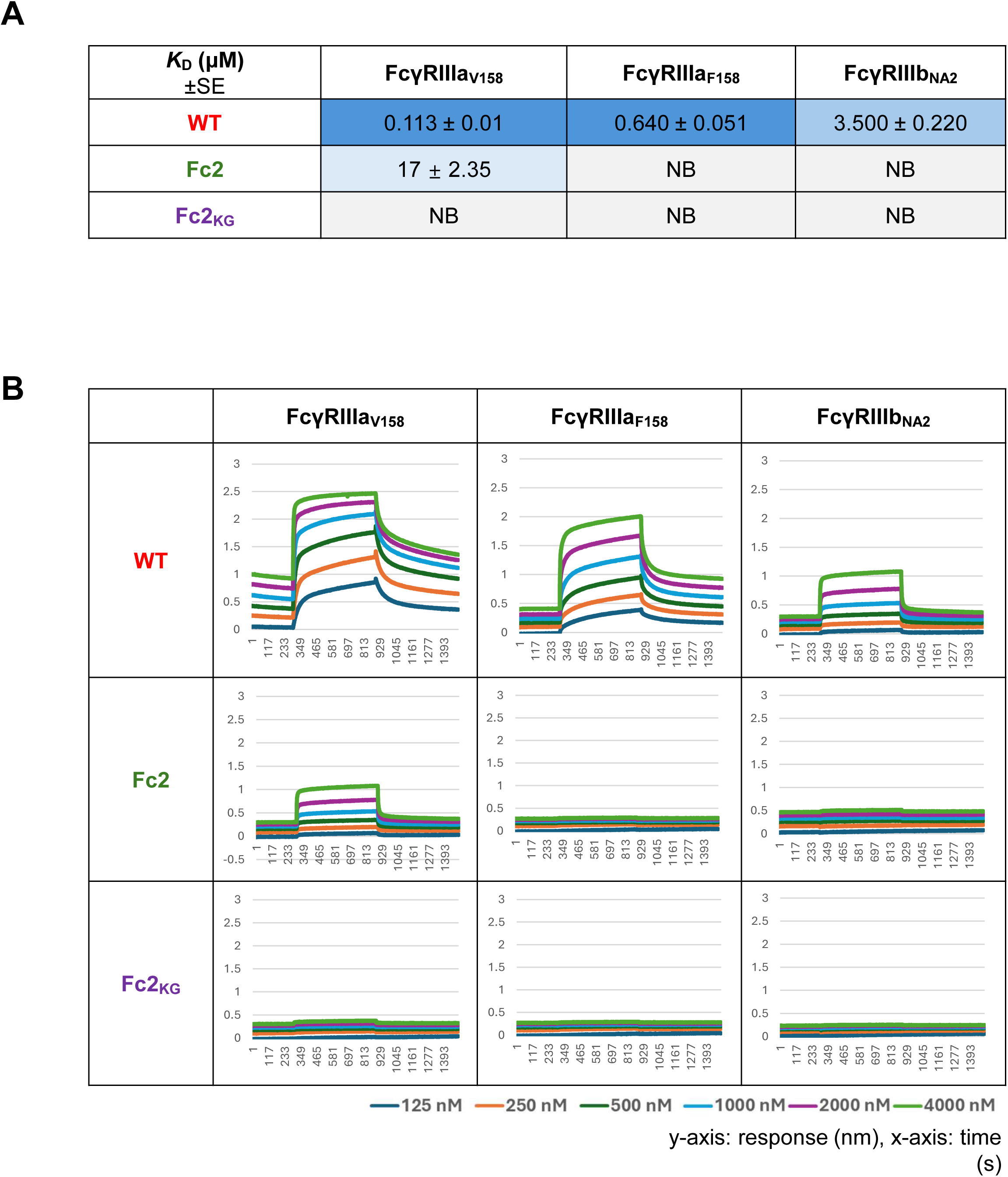

**Figure S2.**
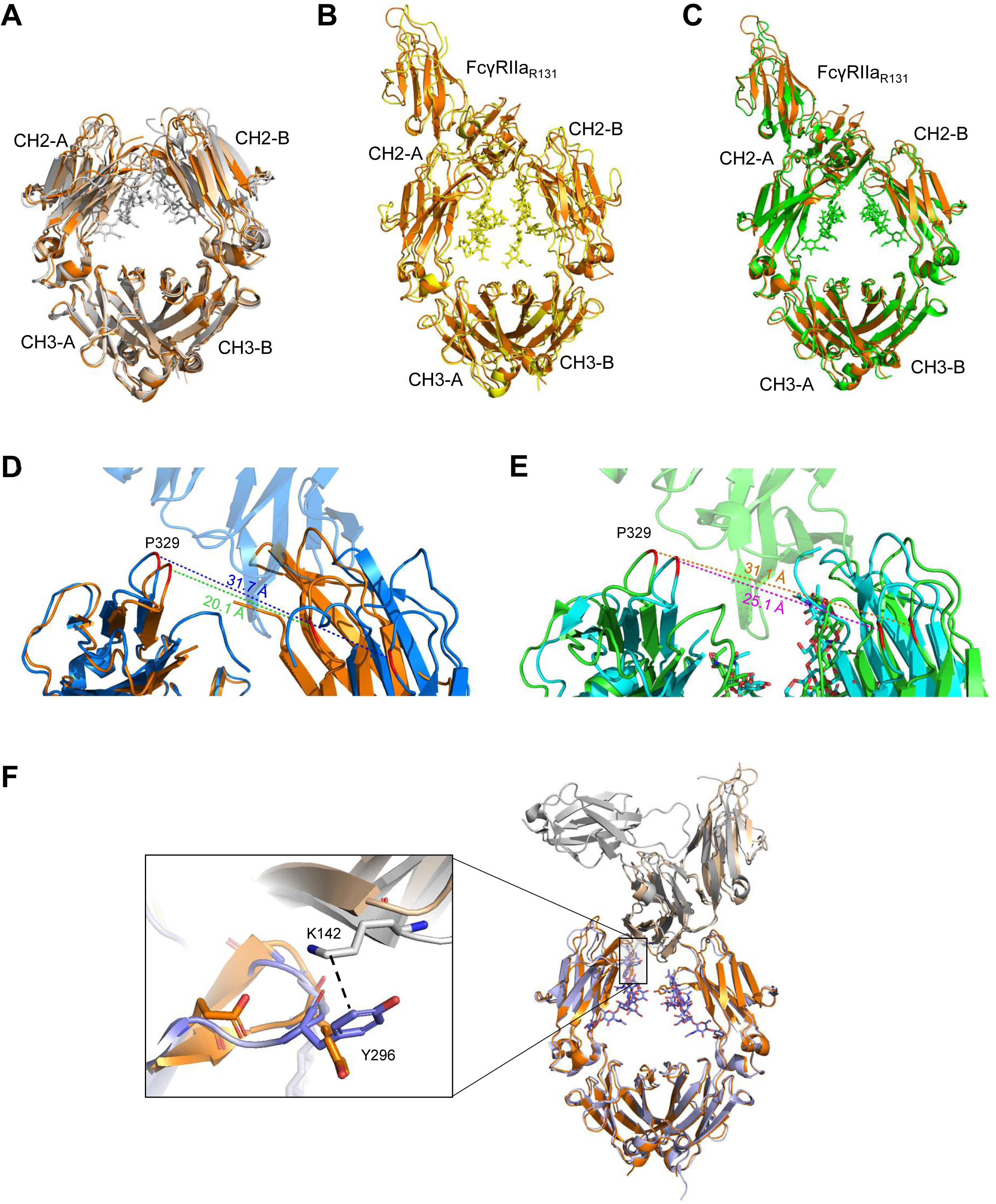

**Figure S3.**
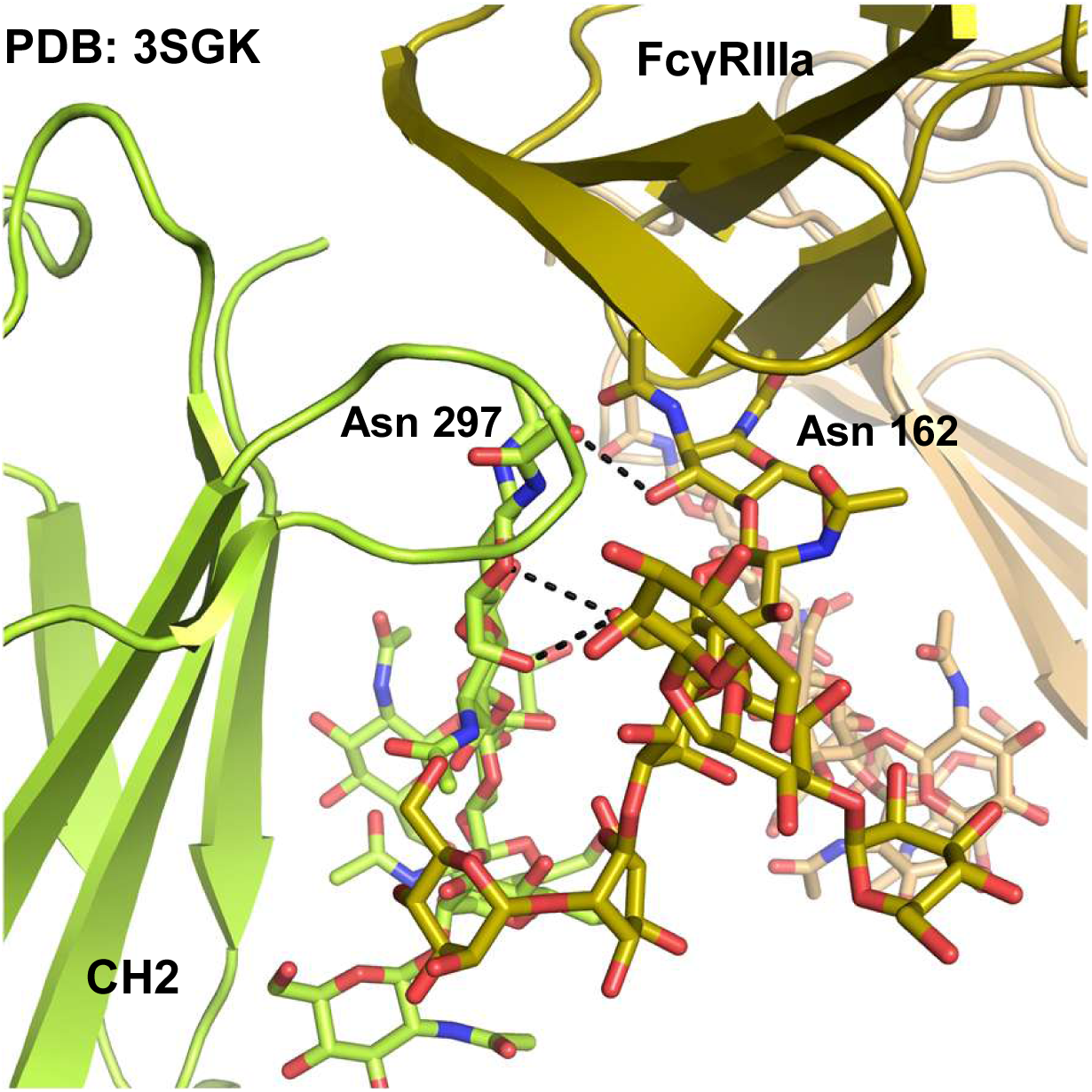

**Figure S4.**
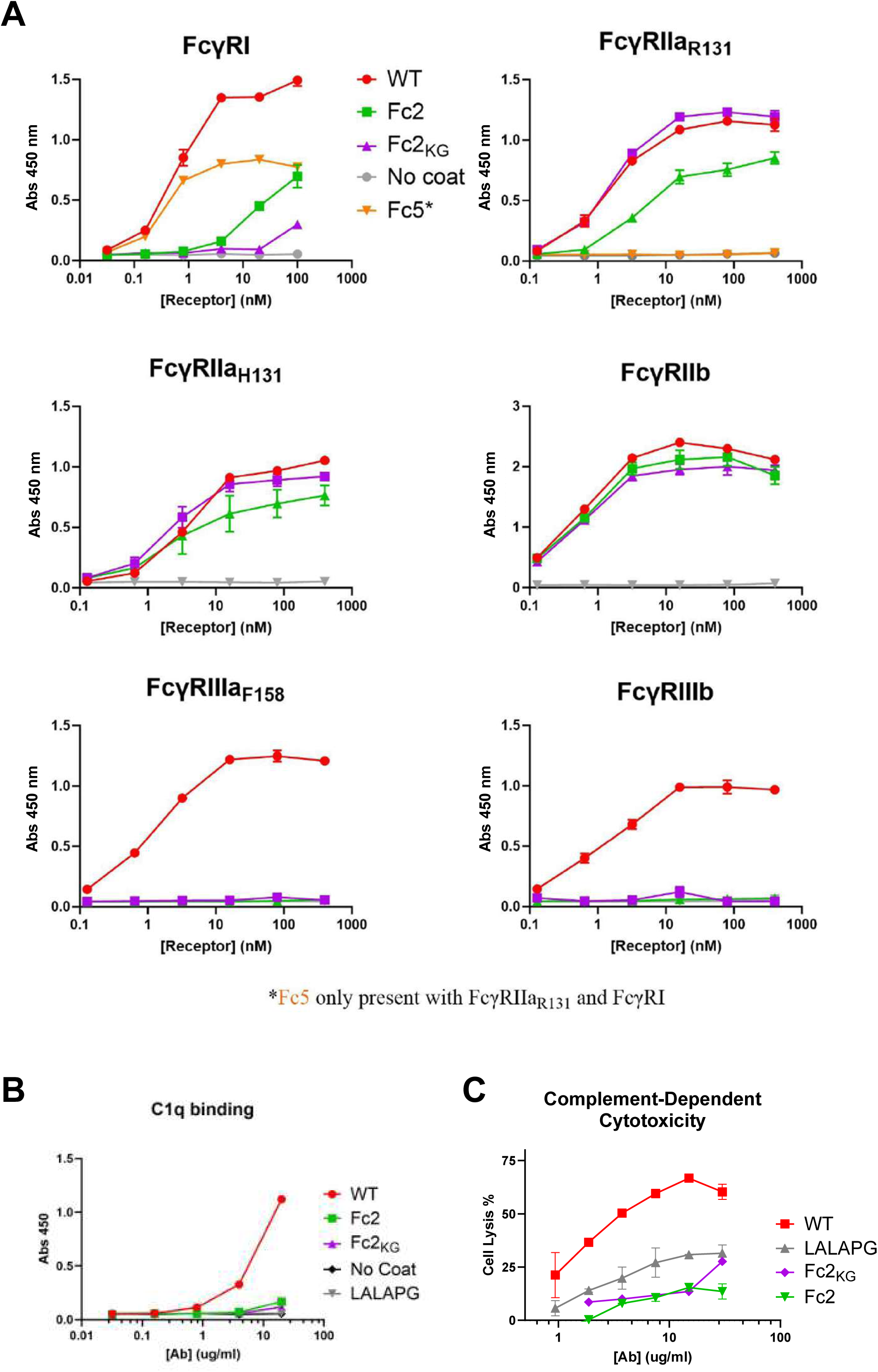

**Figure S5.**
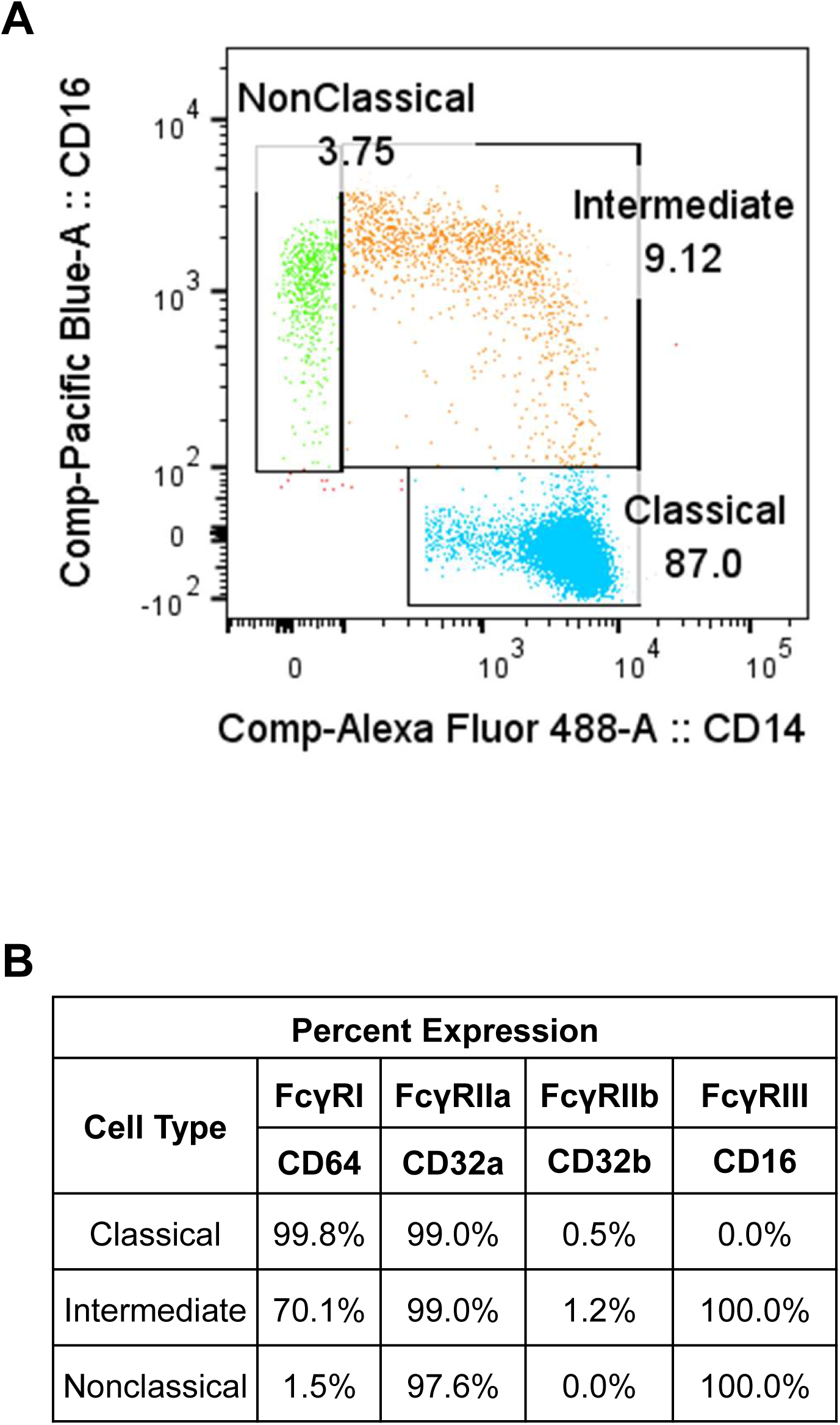

**Figure S6.**
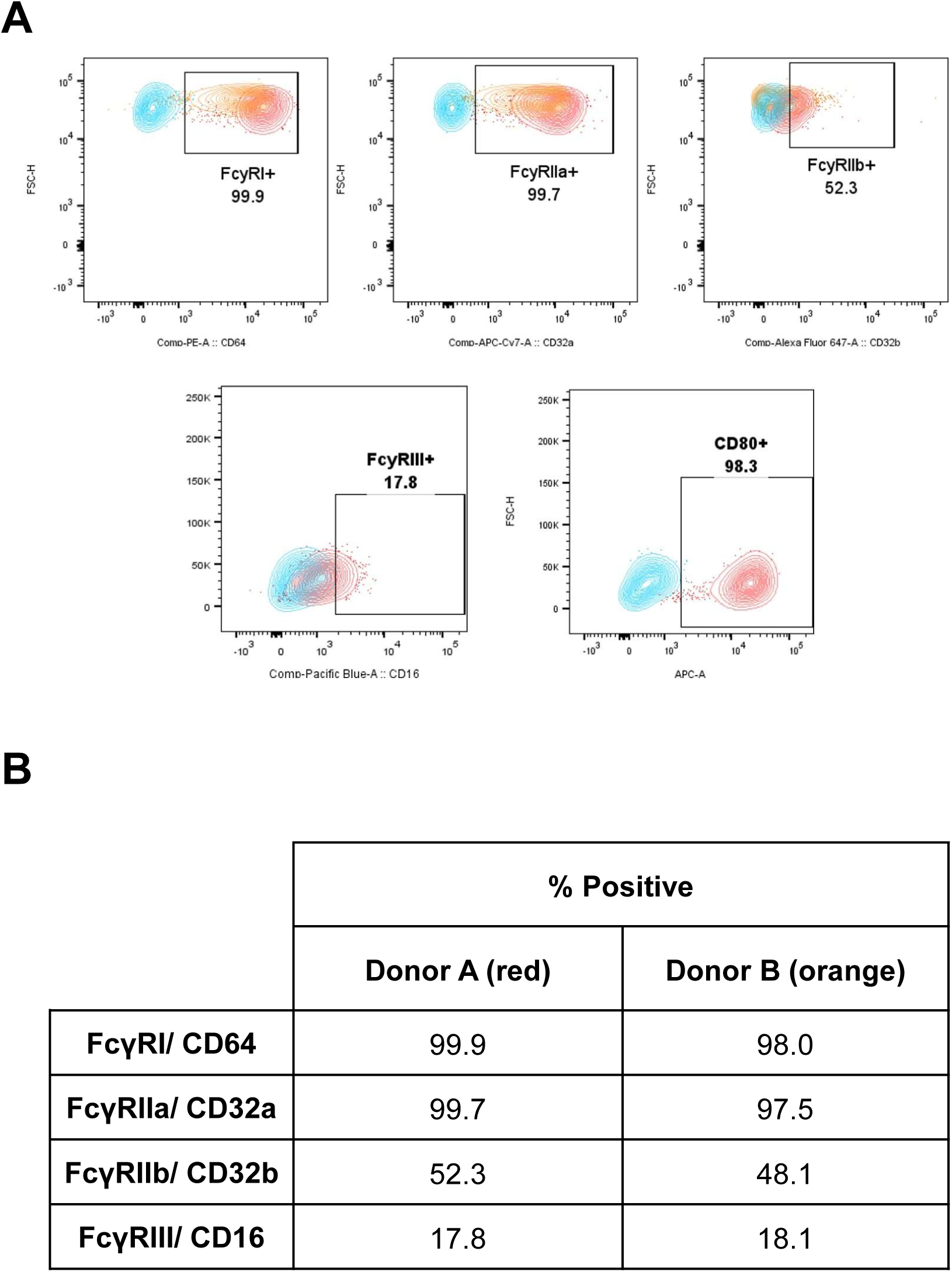

**Table S1.**

| Variant | Mutations |
| --- | --- |
| Fc2 | K246Q, T260A, S298G, T299A, A378N, N390D |
| Fc2<sub>KG</sub> | G236A, K246Q, T260A, S298G, T299A, A378N, N390D |
| Fc5 | T299L, E382V, M428I |
| LALAPG | L234A, L235A, P329G |

**Table S2.**
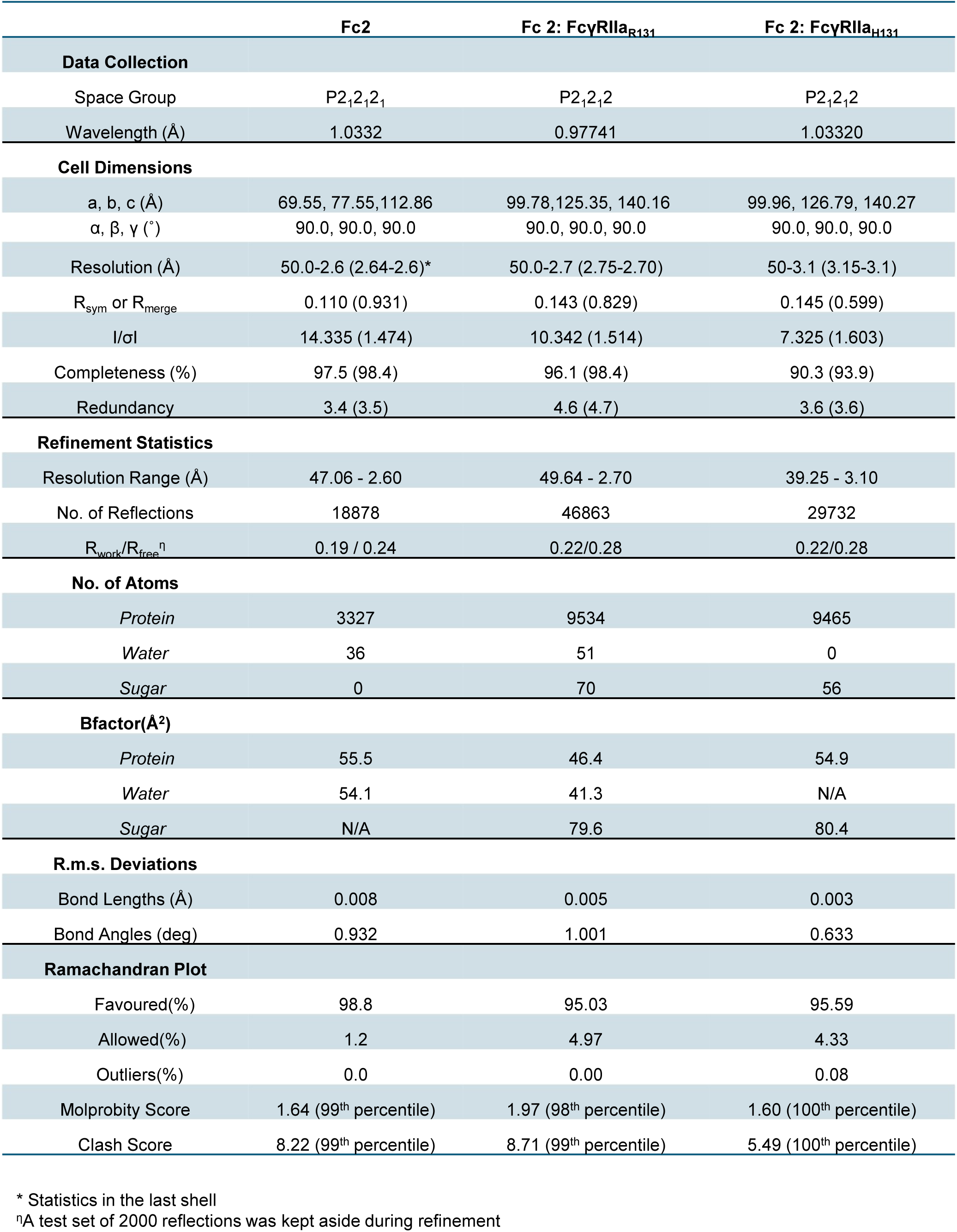

