## Supplementary material for "Antibodies with Engineered Fc Domains Having Absolute Binding Selectivity to either FcγRIIa or FcγRI Delineate the Respective Effector Phenotypes by Human Monocytes and Macrophages": SI figure

**SUPPLEMENTAL INFORMATION**

**Table S1. Engineered IgG1 Fc variants and corresponding amino acid mutations.**

**Table S2. Crystallization parameters for apo Fc2 and co-complex structures of Fc2 with FcγRIIa$_{R131}$ and FcγRIIa$_{H131}$.**

**Figure S1. BLI analysis of engineered IgG1 Fc variants binding to Fc receptors FcγRIIIa$_{V158}$, FcγRIIIa$_{F158}$, and FcγRIIIb$_{NA2}$. A**) Summary of $K_D$ values for rituximab-wt, rituximab-Fc2, and rituximab-Fc2$_{KG}$. All values were calculated using steady-state analysis. **B**) Representative BLI sensorgrams of rituximab-wt, rituximab-Fc2, and rituximab-Fc2$_{KG}$ binding to FcγRIIIa$_{V158}$, FcγRIIIa$_{F158}$, and FcγRIIIb$_{NA2}$. Biosensors immobilized with the indicated Fcγ receptors were dipped into serial concentrations of IgG (125–4000 nM) for association, followed by dipping into buffer alone for dissociation. x-axis: time (s); y-axis: response (nm). Colors correspond to IgG concentrations as indicated.

**Figure S2. Structural characterization of Fc2 and conformational changes upon FcγRIIa engagement. A)** Superposition of wild-type glycosylated IgG1 Fc (gray, PDB: 3AVE), wild-type aglycosylated IgG1 Fc (beige, PDB: 3S7G), and apo Fc2 (orange). **B)** Superposition of wild-type glycosylated IgG1 Fc:FcγRIIa$_{R131}$ (yellow, PDB: 3RY6) and Fc2:FcγRIIa$_{R131}$ (orange). **C)** Superposition of wild-type glycosylated IgG1 Fc:FcγRIIa$_{R131}$ (green, PDB: 9MCY) and Fc2:FcγRIIa$_{R131}$ (orange). **D)** Superposition of apo Fc2 (orange) and Fc2:FcγRIIa$_{R131}$ complex (blue). The Pro329 Cα–Cα distance between the A and B chains increases from 20.1 to 31.7 Å upon receptor binding, reflecting an outward splaying of the two CH2 domains. **E)** Superposition of wild-type glycosylated IgG1 Fc before (cyan, PDB: 3AVE) and after (green, PDB: 9MCY) binding to FcγRIIa$_{R131}$. The distance between the P329 of the A and B chains of the Fc domain is highlighted to illustrate conformational changes upon receptor engagement. **F)** Proposed structural basis for the reduced binding of Fc2 to FcγRI. Close-up view of the Fc2:FcγRIIa$_{R131}$ complex (orange/beige) superimposed on the wild-type IgG1 Fc:FcγRI complex structure (PDB: 4W4O; blue/gray). Zoomed-in box: FcγRI residue K142 and IgG1 Fc residue Y296 observed in the wild-type IgG1 Fc:FcγRI complex (dashed line) and in the Fc2:FcγRIIa$_{R131}$ complex.

**Figure S3. Glycan-mediated contacts at the IgG1 Fc–FcγRIIIa interface.** Close-up view of the binding interface between wild-type glycosylated IgG1 Fc (green, CH2-A chain) and FcγRIIIa (gold) derived from the crystal structure PDB: 3SGK. The Asn297-linked glycan of the IgG1 Fc (green sticks) and the Asn162-linked glycan of FcγRIIIa (gold sticks) form direct glycan–glycan contacts (dashed lines).

**Figure S4. Fc receptor binding and complement activity of engineered Fc variants. A)** Immobilized antibodies were tested for binding to serial dilutions of FcγRII- and FcγRIII-tetramers or FcγRI monomer by ELISA. Representative data shown of one out of two independent experiments run in duplicate. **B**) Binding of serially diluted C1q to immobilized rituximab-wt or rituximab-Fc variants was evaluated by ELISA. **C**) Complement-dependent cytotoxicity (CDC) of Ramos cells opsonized with serially diluted rituximab-wt, rituximab-$Fc2_{KG}$, rituximab-Fc5, or rituximab-LALAPG, followed by incubation with 10% pooled human serum as a complement source. Data shown are representative of two independent experiments performed in duplicate at each concentration. Error bars indicate ± SD.

**Figure S5. Flow cytometric characterization of FcγR expression on primary human monocytes. A)** The expression of FcγRs on classical ($CD14^{++}$ $CD16^{-}$), intermediate ($CD14^{++}$ $CD16^{+}$) and non-classical monocytes ($CD14^{low}$ $CD16^{++}$) was determined by flow cytometry. **B)** FcγRs expression on blood-derived primary monocytes as determined via flow cytometry. The data shown are from one representative of three experiments.

**Figure S6. Flow cytometric analysis of FcγR expression on monocyte-derived M1 macrophages. A)** Representative contour plots showing FcγR surface expression on monocyte-derived M(LPS+IFNγ) macrophages from two donors after 9 days of in vitro differentiation and polarization. The blue population represents unstained cells from Donor A, the red population represents Donor A stained with the indicated fluorophore-conjugated antibody, and the orange population represents Donor B stained with the indicated fluorophore-conjugated antibody. CD80 expression was included as a marker of M1 polarization. **B)** Quantitative summary of FcγR surface expression (% positive) on M(LPS+IFNγ) macrophages from two independently differentiated donors. Colors correspond to those in panel (A): Donor A (red) and Donor B (orange). Gating was performed as described in Figure S6A.
